# A Simulator for Somatic Evolution Study Design

**DOI:** 10.1101/2022.05.01.487551

**Authors:** Arjun Srivatsa, Haoyun Lei, Russell Schwartz

## Abstract

**Motivation:** Somatic evolution plays a key role in development, cell differentiation, and normal aging, but also diseases such as cancer, which is now mainly thought of as a disease of genetic and epigenetic modification. Understanding mechanisms of somatic mutability — variant types and frequencies, phylogenetic structure, mutational signatures, and clonal heterogeneity — and how they can vary between cell lineages will likely play a crucial role in biological discovery and medical applications. This need has led to a proliferation of new technologies for profiling single-cell variation, each with distinctive capabilities and limitations that can be leveraged alone or in combination with other technologies. The enormous space of options for assaying somatic variation, however, presents unsolved informatics problems with regards to selecting optimal combinations of technologies for designing appropriate studies for any particular scientific questions. Versatile simulation tools are needed to make it possible to explore and optimize potential study designs if researchers are to deploy multiomic technologies effectively.

**Results:** In this paper, we present a simulator allowing for the generation of synthetic data from a wide range of clonal lineages, variant classes, and sequencing technology choices, intended to provide a platform for effective study design in somatic lineage analysis. Our simulation framework allows for the assessment of study design setups and their statistical validity in determining different ground-truth cancer mechanisms. The user is able to input various properties of the somatic evolutionary system, mutation classes (e.g., single nucleotide polymorphisms, copy number changes, and classes of structural variation), and biotechnology options (e.g., coverage, bulk vs single cell, whole genome vs exome, error rate, number of samples) and can then generate samples of synthetic sequence reads and their corresponding ground-truth parameters for a given study design. We demonstrate the utility of the simulator for testing and optimizing study designs for various experimental queries.

**Availability:** https://github.com/CMUSchwartzLab/MosaicSim

## 1 Introduction

Advanced sequencing technologies have made it possible to profile genetic variation at the single-cell level on population scales, revealing in part that the human body is a continuously evolving genetic mosaic (11; 46; 1). Genetic and epigenetic modifications in somatic cells over many generations of cell growth and replication result in heterogeneity between cells, tissues, and organs in normal aging and development. Somatic mosaicism can also produce disease phenotypes, such as neurodegeneration and, most notably, cancer (29; 35). Profiling somatic mutations is an active area of research for biological discovery. It is also rapidly becoming a standard practice in cancer care, where clinicians now commonly utilize gene panels to screen for common “driver” mutations (43; 18). Accumulating genomic data has made it apparent that somatic mutability is much more complicated than early models of tumor clonal evolution first suggested (5) and far more extensive in even healthy tissues (c.f.,(4; 7)). Somatic variation produces complex patterns of “mutational signatures” (14; 2; 21) reflecting different endogenous (e.g., DNA replication processes) and exogenous (e.g., UV light) mechanisms of mutability. In cancers and other precancerous conditions, high levels of somatic mutability are frequently observed due to damage to cell replication on error-correction machinery (28; 27; 39). They further may include not just single nucleotide variations (SNVs) but potentially extensive copy number alterations (CNAs) and structural varations (SVs), including complex mutation events rarely seen in conventional population genetics, such as chromoplexy and chromothripsis (26).

As we have come to understand the extent and importance of somatic evolution, enormous effort has been put into developing biotechnological tools for profiling somatic variability at ever greater scales, precision, and accuracy (8; 40). No one technology is able to comprehensively characterize somatic variability across a complex tissue and do so with precision and accuracy and at low cost. Rather, investigators attempting to characterize somatic variation processes now have available to them a vast array of technologies — e.g., short read vs. long read vs. single-cell sequencing, liquid biopsy vs. tissue biopsy, whole genome vs. whole exome vs. targeted sequencing — each with distinctive different properties and tradeoffs (42). Current work increasingly depends numerous possible multiomic data combinations (e.g., long-read and short-read or bulk and single-cell data), along with various other study design choices (e.g., number of replicates, depth of coverage), with uncertain knowledge of how these choices together with different choices of analysis software will influence one’s ability to quantify any particular feature of the somatic evolution process (32; 20). There is currently little empirical or theoretical basis on which an investigator planning a study can select a combination of technologies and study design well suited for any particular investigation.

Simulation presents a viable solution to these issues by allowing for efficient tests of various study designs with direct knowledge of most biological parameters of interest. Various sequencing simulators have been developed that make it possible in principle to test a study design before the potentially large expense of executing it in the lab. For example, the popular BAMSurgeon simulator (10; 23) imposes variation onto existing BAM alignment files for realistic tumor reads. Most such simulators focus on generating accurate reads from existing sequencing technologies and companies (9; 49). While these simulators are useful in benchmarking tools, they are not generally designed for the purpose of considering different study designs, particular ones that involve hybrid sequencing modalities or broad coverage of variant types. To our knowledge, no existing simulator allows for the testing of broad forms of genetic variation, sampling strategies, and hypothetical sequencing properties. Most sequence simulators to date are targeted towards population genetics evolutionary problems and fail to capture the broad ranges of mutation mechanisms and rates observed in somatic evolution, particularly in hypermutable conditions frequently seen in cancers. The simulators created specifically for cancerous conditions, on the other hand, tend to focus on one specific technology or aspect of the cancerous condition. For example, Mallory et al. (30) designs a simulator specifically for copy number analysis in single-cell sequenced data, and Nicol et. al (33) focus primarily on the spatial distribution of mutations.

Here, we seek to meet the needs of sequencing study design for somatic variation studies through a new clonal evolution simulator. Our simulator links a coalescent model of clonal evolution to a versatile model of read generation with user-configurable variant classes, mutation rates, evolutionary models, sequencing setups, and study design decisions. Our framework focuses on general properties of sequencing that allow for the design of better experiments and future sequencing technologies. It also introduces a wide variety of features important to somatic variation studies that are not, to our knowledge, found in any other current simulator, such as capturing broad classes of variation like breakage-fusion-bridge, chromothripsis, and chromoplexy that have been implicated in certain cancers (45). A detailed feature comparison between our simulator and related simulators is shown in Table 1 (47; 31; 36; 10). We demonstrate utility of this simulator through application to a series of hypothetical questions in testing and optimizing study design for somatic evolution studies.

**Table 1:** A feature comparison between our simulators and a few other sequencing simulators with related functions.

|  | Our Simulator | pSITE | CellCoal | BAMSurgeon |
| --- | --- | --- | --- | --- |
| Lineage Properties | ✓ | ✓ | ✓ | ✗ |
| Single Cell | ✓ | ✓ | ✓ | ✗ |
| Mutational Signatures | ✓ | ✗ | ✓ | ✗ |
| Variable Read Length | ✓ | ✗ | ✗ | ✗ |
| Variable Error Rate | ✓ | ✗ | ✗ | ✗ |
| SNP/Indel Variation | ✓ | ✓ | ✓ | ✓ |
| Resource Usage | Medium | Medium | Low | Medium |
| Multiple Sites/Samples | ✓ | ✗ | ✓ | ✗ |
| Runtime length | Medium | High | Low | Medium |
| Copy Number Variation | ✓ | ✓ | ✗ | ✓ |
| Rare Structural Variation | ✓ | ✗ | ✗ | ✓ |
| Whole Genome Reads | ✓ | ✓ | ✗ | ✓ |
| Whole Exome Reads | ✓ | ✓ | ✗ | ✓ |
| Targeted Sequencing Reads | ✓ | ✗ | ✗ | ✓ |
| Mutation Frequencies | ✓ | ✓ | ✓ | ✗ |
| Mutation Distributions | ✓ | ✗ | ✓ | ✗ |
| Clonal Frequency Parameters | ✓ | ✗ | ✓ | ✗ |
| Sequencer Quality Consideration | ✗ | ✓ | ✓ | ✓ |
| RNA Sequencing | ✗ | ✗ | ✗ | ✓ |
| Liquid Biopsy | ✓ | ✗ | ✗ | ✗ |

## 2 Materials and Methods

The complete simulator consists of four main modules: 1) sampling an evolutionary lineage tree for the clonal evolution, 2) sampling mutation events on the lineage tree, 3) simulating sequence reads, and 4) sampling reads based on experimental design decisions. Below, we describe each module in turn. Each module has a number of user-tunable parameters to control different biological parameters of the presumed cell lineages as well as experimental parameters of the sequencing strategy. The main tunable parameters of the simulator are summarized in Table 2. We describe the individual modules in more detail in the subsequent subsections. Full pseudocode for these modules can be found in the appendix.

**Table 2:**
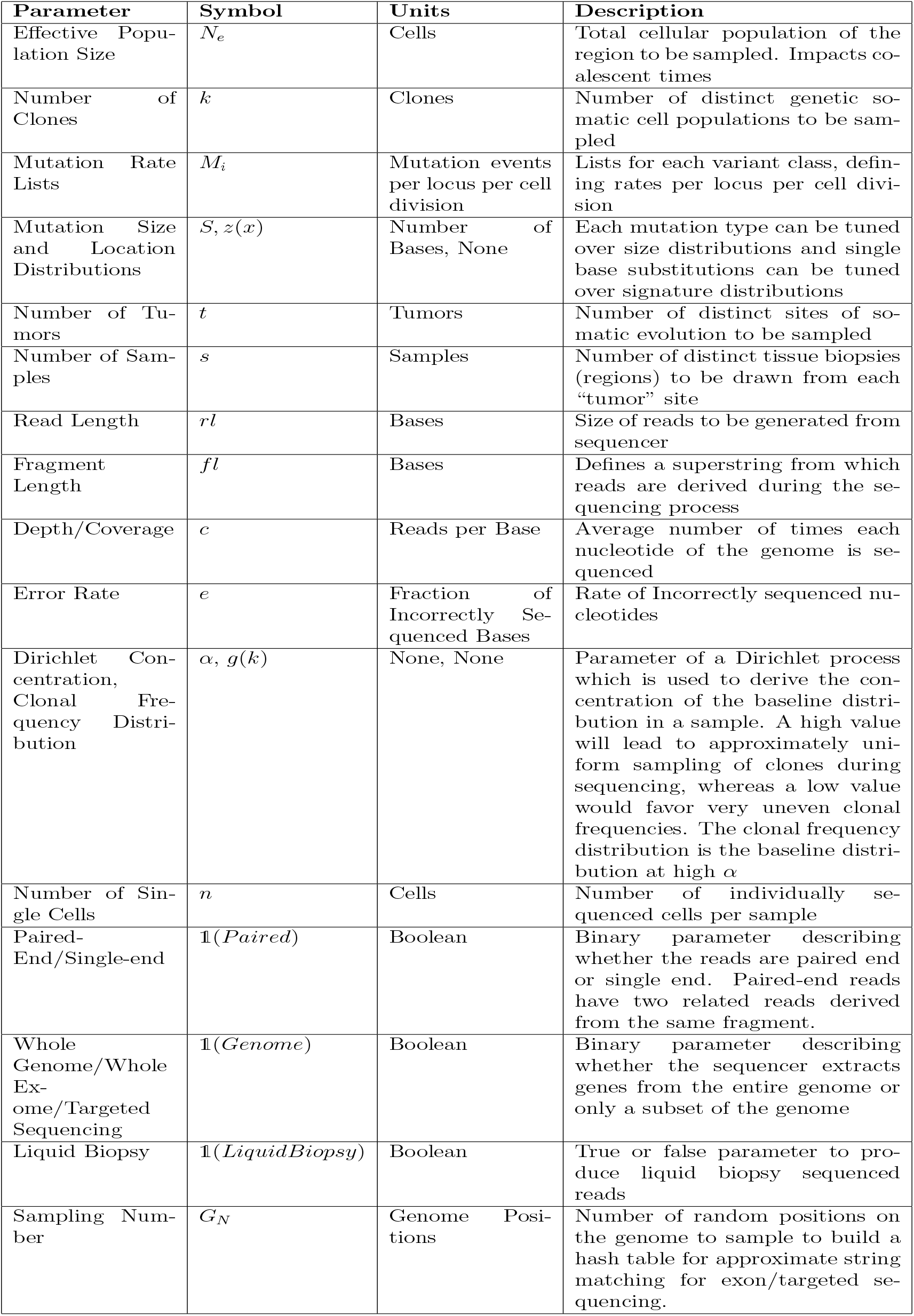
Summary of the main tunable parameters

| Parameter | Symbol | Units | Description |
| --- | --- | --- | --- |
| Effective Population Size | $N_e$ | Cells | Total cellular population of the region to be sampled. Impacts coalescent times |
| Number of Clones | $k$ | Clones | Number of distinct genetic somatic cell populations to be sampled |
| Mutation Rate Lists | $M_i$ | Mutation events per locus per cell division | Lists for each variant class, defining rates per locus per cell division |
| Mutation Size and Location Distributions | $S, z(x)$ | Number of Bases, None | Each mutation type can be tuned over size distributions and single base substitutions can be tuned over signature distributions |
| Number of Tumors | $t$ | Tumors | Number of distinct sites of somatic evolution to be sampled |
| Number of Samples | $s$ | Samples | Number of distinct tissue biopsies (regions) to be drawn from each “tumor” site |
| Read Length | $rl$ | Bases | Size of reads to be generated from sequencer |
| Fragment Length | $fl$ | Bases | Defines a superstring from which reads are derived during the sequencing process |
| Depth/Coverage | $c$ | Reads per Base | Average number of times each nucleotide of the genome is sequenced |
| Error Rate | $e$ | Fraction of Incorrectly Sequenced Bases | Rate of Incorrectly sequenced nucleotides |
| Dirichlet Concentration, Clonal Frequency Distribution | $\alpha, g(k)$ | None, None | Parameter of a Dirichlet process which is used to derive the concentration of the baseline distribution in a sample. A high value will lead to approximately uniform sampling of clones during sequencing, whereas a low value would favor very uneven clonal frequencies. The clonal frequency distribution is the baseline distribution at high $\alpha$ |
| Number of Single Cells | $n$ | Cells | Number of individually sequenced cells per sample |
| Paired-End/Single-end | $\mathbb{1}(Paired)$ | Boolean | Binary parameter describing whether the reads are paired end or single end. Paired-end reads have two related reads derived from the same fragment. |
| Whole Genome/Whole Exome/Targeted Sequencing | $\mathbb{1}(Genome)$ | Boolean | Binary parameter describing whether the sequencer extracts genes from the entire genome or only a subset of the genome |
| Liquid Biopsy | $\mathbb{1}(LiquidBiopsy)$ | Boolean | True or false parameter to produce liquid biopsy sequenced reads |
| Sampling Number | $G_N$ | Genome Positions | Number of random positions on the genome to sample to build a hash table for approximate string matching for exon/targeted sequencing. |

### 2.1 Lineage Simulation

For each simulation, we generate a cell lineage assuming that mutations are selectively neutral and generally follow the assumptions of the standard coalescent model (34). User-definable parameters include a total population size (*N*_*e*_), set to 8 × 10^8^ cells by default, as well as a number of clones (*k*) to be sampled. Coalescent sampling of tree topologies and edge lengths is implemented using msprime (17). The unit of time we use is a generation, i.e., the time of a single cell division.

### 2.2 Mutation Events

The simulator supports commonly discovered types of somatic variations, particularly those implicated in the development of cancer (44; 6). These currently include the following: single nucleotide variation (SNVs), copy number abberations (CNAs), insertions, deletions, kataegis, chromothripsis, chromoplexy, aneuploidy, translocations, inversions, and breakage fusion bridge cycles (BFBs). Table 3 summarizes these mutation types. Each mutation type is implemented by sampling from various probability distributions for location and length while simultaneously maintaining constraints encapsulating our knowledge of the mechanism for each mutation type. The scale of many of these forms of variations can be tuned, but with default values set based on estimated distributions of sizes found in current studies of cancer genomes (26). Size distributions for structural variants are modeled as a truncated mixture of negative binomials to represent small, medium, and large scale events. More detail on implementations for these mutation types are listed in the supplementary materials. Each mutation type also has a rate at which that mutation appears which we assume may differ between tumor stages and in healthy tissues per the “mutator phenotype” hypothesis (27). The simulator also currently supports simulating mutations drawn from single base substitution signatures derived from the COSMIC dataset. As more signatures are discovered and validated, these can be readily incorporated into the current mutation framework. Similarly, distributions over mutation location and size are flexible.

**Table 3:** Description of Available Mutation Types

| **Mutation Type** | **Description** |
| --- | --- |
| Single Nucleotide Variation(SNV) | A single base is changed to a different nucleotide |
| Single Base Signature | A single base is mutated with frequency according to its trinucleotide context, termed a mutational signature |
| Copy Number Variation(CNV) | A random region of the genome is repeated a number of times |
| Inversion | A random region of the genome has its nucleotide sequence reversed |
| Deletion | A random region of the genome is removed |
| Translocation | A trailing portion of one chromosomal sequence is either added or exchanged with a trailing part of another chromosome sequence |
| Insertion | A random sequence of a given length is inserted at a random position in the genome |
| Aneuploidy | A chromosome is either deleted or copied a random number of times |
| Kataegis | Within a randomly sampled region of the genome, a particular type of single base substitution (e.g., C>T) occurs with high frequency |
| Chromothripsis | A region of a single chromosome is broken into many pieces, these pieces are deleted with a certain frequency, and finally rearranged randomly to replace the selection region. |
| Chromoplexy | Multiple chromosomes are shattered into pieces and rearranged in an order consistent with telomere endings |
| Breakage Fusion Bridge | A chromosome loses a telomeric segment and proceeds to repeatedly break and fuse with itself to create new chromosomes |

Once a lineage is simulated, we apply mutations to this lineage going forward from the most recent common ancestor of all the clones to be sampled. Mutation rates for each class of variation are defined as a uniform discrete distribution of potential rates, *M*_*i*_. For each edge of the lineage, a specific rate is generated via *r*_*i*_ ∼ *M*_*i*_ and mutation times are generated via a Poisson process, with rate *r*_*i*_. This is done for each edge of the phylogeny independently for each class of mutation. The end result of this process is a list of times for each mutation type and its occurrence on each edge of the phylogeny. Next, the simulator imposes each of these mutations on a reference genome to establish the sequences of clones at all nodes of the lineage tree. Given that mutations were independently simulated for each mutational class, we first merge and sort the mutational events. For a user selection of *k* for the total number of clones, there are 2*k* − 1 total nodes, including the root node. We first use the tree data-structure to compute all root-to-leaf paths in the tree, which allows us to generate all potential clonal genomes. We start with the root node as the reference and impose sorted mutations for each root-to-leaf path. Once an internal node has been computed, it is never recomputed so as to save computation. The end result of this process is a stored genomic sequence for each clone, including those at internal nodes, that is later sampled in the sequencing step.

### 2.3 Sequencing Implementation

Sequencing procedures differ depending on the type of sequencing chosen (e.g., whole genome sequencing or targeted sequencing). The general strategy, however, is similar. First, a clone from the tree is sampled from a Dirichlet process, i.e. *k* ∼ *G* ∼ *DP*(*g*(*k*), *α*). The sampled clone may be either a leaf node or an internal node, but not the root node. For this selected clone, stochastic fragment lengths are drawn following *fl* ∼ *TruncatedNegBin*(*rl*). That is, we draw samples from a negative binomial with expectation *rl*, and discard fragments with length below 30. Using these fragment lengths, operations are performed on fragmented clonal reads defined by the user-parameters. Specifically, clone *k* is loaded, “chopped” according to *fl*, subsetted based on *rl* and 𝟙(*Paired*), seeded with errors according to *e*, and written to a FASTQ file. After this particular clone is sequenced, the coverage of the simulator is updated according to the fraction of the genome covered depending on the read and fragment length. This process repeats with independent repeats of each stochastic process until coverage *c* is reached.

The process for exome and targeted sequencing requires additionally identifying reads that align to exon or targeted gene sequences. This approximate string matching problem is computationally infeasible over every possible read so we make a few simplifying assumptions. First, we create k-mer sets for each of our target sequences and use locality sensitive min-hashing to index these sequences for fast lookup (41; 37). Then, for genome locations surrounding our target sequence intervals, we calculate whether reads originating from these genome locations match some target sequence in the hashing index above a certain threshold. If this location matches some sequence, then we add it to a list of locations. We repeat this sampling *G*_*N*_ times, and use this list combined with the original sequence locations to generate a discrete probability density to sample read locations. Without large scale disruptions to genome structure, the user may opt to set *G*_*N*_ to zero, implying that the original exon locations serve as good approximations for read matches. We sample reads from this generated list with the given read length while simultaneously sampling different clones until our desired coverage is reached. Targeted sequencing is performed in much the same way as exon sequencing, but the target sequences are defined by a different text file of chromosome, start, and end locations.

Modifications also occur for single cell sequencing, where we do not continually re-sample cell clones, but instead sample only once and use the given clone until the desired coverage is reached. Finally, for liquid biopsy, we perform a similar iterative clonal sampling procedure, but do not chop the sequence uniformly. Rather, we draw random fragments from the genome strings of each clone and mix them at some frequency with reference DNA then return these in a read file.

### 2.4 Experimental Decisions

The full simulation is defined by looping the three modules — lineage generation, mutation sampling, and read generation — over tumors and samples. Specifically, we independently define and execute a separate lineage sampling and mutational frequency sampling *t* times, one for each tumor. Similarly, we configure parameters related to sequencing decisions (*rl*/*fl, c,n*, 1(*Paired*), etc.), and execute the procedures listed in Section 2.3, *s* times per tumor. A note with respect to single-cell sequencing is that a number of single cells is defined for a single sample, and a subdirectory is created for that sample. By default, we assume that each run of the simulator creates a “reference” sample, which is simply a single-cell sample on the root node of the tree. The final result of the simulator is a labeled directory with sub-directories corresponding to reference reads, tumor reads, and sample reads as depicted in Figure 1. Each of these directories holds ground truth parameters with information about the tumor and sequencing parameters.

**Figure 1:**
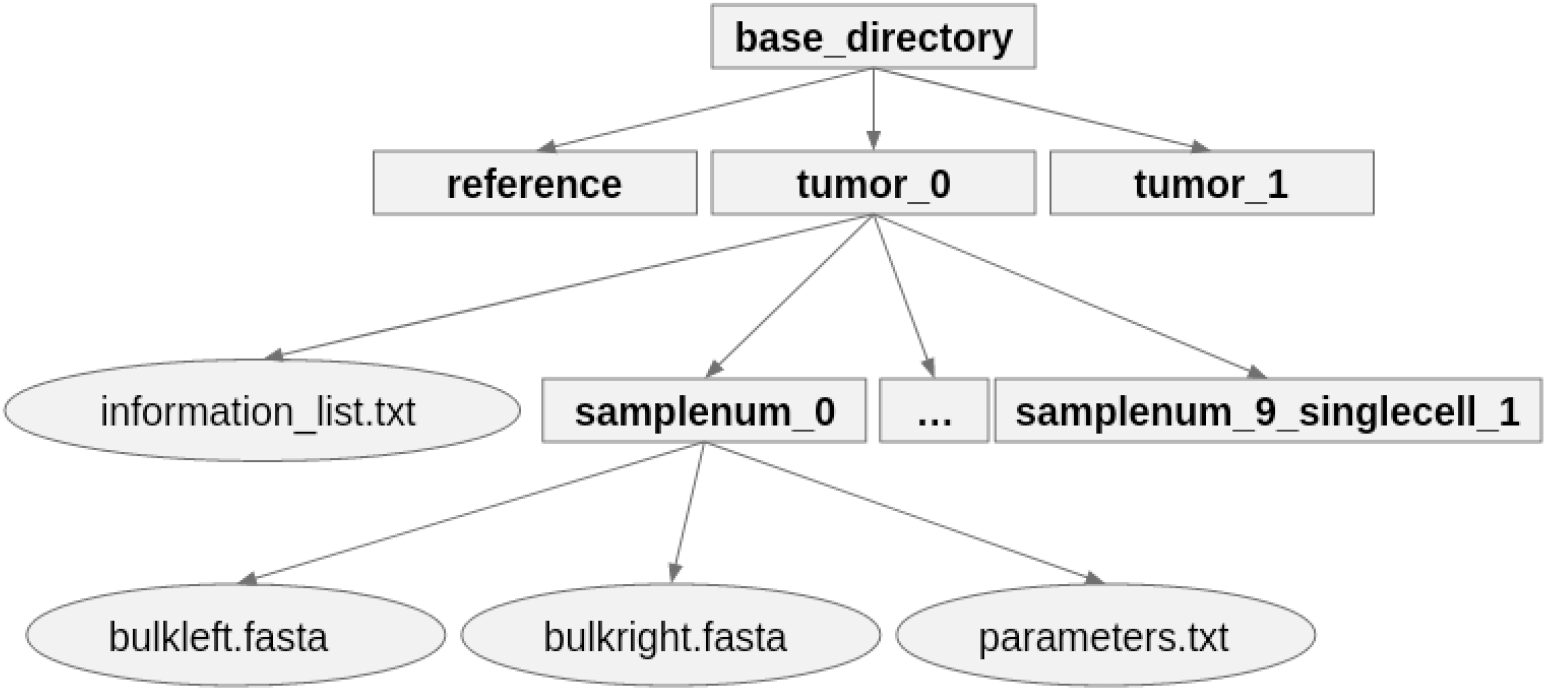
Sample computer output of the simulator is shown above. Directories are depicted as boxed text, whereas files are shown in ovals.

While we described the simulation method for singular values of the parameters listed in Table 2, the practical implementation of the simulator encodes most of the parameters as a list of values and samples from each list to generate the total simulation. This random sampling procedure allows for grid search exploration of study design spaces more easily. We refer to each parameter with a subscript to denote the sample number. For example, *n*_*i*_ would denote the number of single cells on the *i*^*th*^ sample. In cases where we survey multiple sites of somatic evolution, we use a double subscript to remove ambiguity. For example, *c*_*ij*_ would denote the coverage of the *j*^*th*^ sample on the *i*^*th*^ tumor.

### 2.5 Runtime and Space Analysis

The runtime of the program is generally dominated by the sequencing steps rather than the coalescent or mutation steps. Defining *c*_*ij*_ as the sequencing coverage of the *j*^*th*^ sample on tumor *t*_*i*_, *n*_*ij*_ as the number of single cell samples drawn on the on the *j*^*th*^ sample of tumor *t*_*i*_, and *s*_*i*_ as the number of samples drawn on tumor *t*_*i*_. The run time is approximately proportional in 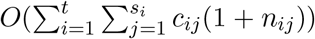, which approximately calculates the amount of times we traverse and sequence the genome.

The maximum memory of the program is constrained by the batch and subblock size as well as read length and sequencing type. At its minimum value of one and with standard read sizes in the range of 75 to 1000 base pairs, the maximum memory is a small constant factor (generally around 5-10x) of the largest stored clonal genome. If the user wants to minimize memory footprint, then he or she should set the batch and subblock size to 1, and/or avoid generating long-read exome sequenced data.

The amount of non-volatile (disk) storage of the program is bounded by the storage of the clonal genomes during the mutational process as well as the sequencing FASTQ files. This is approximately proportional to 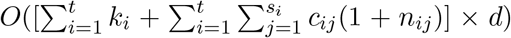, where *d* is the genome size, *k*_*i*_ denotes the number of clones in tumor *t*_*i*_ and *c*_*ij*_ and *n*_*ij*_ are defined as above. Many of the files are intermediary and can be removed after post-processing, if desired.

### 2.6 Computational Implementation and Resource Requirements

Code for the simulator is available in Python, with additional analysis code in Python scripts. Executing one run of the simulator involves changing/setting the parameters and running the python command. Multiple simulations can be performed in parallel on separate threads, provided the system has enough memory. Simulations can also be distributed across a multi-node cluster for large scale data generation. All our tests were performed on an multi-node Ubuntu system with 184 cores, 850GB of memory, and 10TB of storage. A single run of the simulator only uses one core. As a reference point, three 30x-paired WGS samples take around 3.5 hours to generate on our system and use around 250-300GB of disk storage and less than 40G of memory.

## 3 Experimental Results

In this section, we demonstrate the utility of our simulator in evaluating or optimizing sequencing study designs for profiling clonal evolution. A primary motivation of this simulator is to plan study designs for evaluating questions about differences in somatic mutability between subsets of samples. The questions for study design are to test whether a difference can be detected between two subsets of samples under a given study design or to find a study design optimizing power to detect such differences. Here, study design might include changes in the types of sequencing applied, the informatics software used to evaluate it, and features such as the number of tumors and tumor sites or regions to be examined. We would then assume that the study is being used to test for differences in biological parameters between subsets of samples. Such biological parameters might include mutation rate differences, presence of rare variations, variation in mutational signatures, phylogeny structures, or clonal frequencies – although we mainly focus on mutation rate variations and variant calling in our present tests.

### 3.1 Notation and Performance Measures

For the analyses presented here, we take the sequencing read outputs from our simulator and perform alignment to the hg38 reference genome, after which we call several forms of variation for analysis. The aligners used were minimap2 (25), Bowtie (22), and bwa-mem (24); the callers used were Strelka (19) and Delly (38). Throughout the tests, we reference our study design, which we formally define as a collection of matrices **X** = {*X*_1_, …, *X*_*t*_}, where *t* denotes the number of tumors. The matrices *X*_*i*_ denote the sequencing decisions taken on tumor *i* and encapsulate every sample. Namely, each matrix *X*_*i*_ is of dimension 7 × *s*_*i*_, where *s*_*i*_ denotes the number of samples taken on tumor *i*. The columns of *X*_*i*_ denote sequencing and informatics choices on a single sample *s*_*i*_. That is, the matrix is of the form:

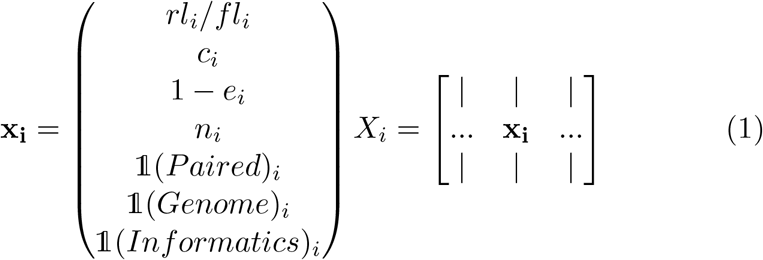

The collection of matrices representing our study design is mostly relevant for an experimenter hoping to test combinations of sequencing and repeated tumor biopsy draws within a patient with multiple neoplastic sites. In our experimental tests, we often consider the single tumor, single sample case. With this instance, we can collapse the collection of matrices down to a single vector **x**.

To judge a study design’s utility, we require performance measures that assess whether the design is recovering a signal related to the intended hypothesis. In our experiments, this often meant assessing the accuracy with which one or more variant call files aligned with the reported set of mutations from the simulation. The problem of assessing a variant call report in a heavily mutated cancer sample is a challenging problem in its own right due to issues of temporality and location consistency. Temporality issues obscure the location of any particular variant, making any final reported variant difficult to trace back to its origin. For example, if a SNV occurred in a region afflicted by a chromothriptic event, the final reference-aligned reported SNV would be at a distinct location from the SNV location on the simulated genome. High fidelity structural variant recovery is even more challenging because the field has not settled on a consistent set of genomic rearrangements, nor ways to reproducibly label these with respect to a linear reference genome. We designed measures derivable from variant call files based on these assumptions. For mutation rate tests, we simply used the raw count of each variant reported in each VCF as an estimate for the amount of each form of variation. The assumption underlying this is that even mislocated or false positive variants are indicative of the overall mutation rate of the sample and relatively consistent across samples. To assess the actual accuracy of SNV tests, we used two primary measures. The first is simply recall, or the proportion of the actual events that we recovered, i.e.: 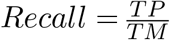, where *TP* defines the count of true positive mutations and *TM* is the total number of ground truth mutations. We favor recall as a measure in highly mutated or error-prone samples, where callers will return a high fraction of false positives. The alternative measure we use is the *F*1 score which provides a more detailed view of the classifier accounting for false positives, that is: 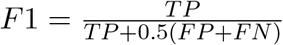, where *TP* stands for the true positive counts, *FP* for false positive counts, and *FN* for false negative counts. Some of the false positive counts may actually be correct variants, but the location mapping with respect to variants on sequentially simulated clonal genomes is difficult to recover. For structural variants, we designed a measure with the assumption that any particular reported event is more likely to be referring to a ground truth event if they overlap. Call the output of the variant calling software for chromosome *i, C*_*i*_ = {(*a, b*)}, and call our ground truth set of structural variant locations, *D*_*i*_ = {(*c, d*)}, where tuples (*a, b*), (*c, d*) represent all starting and ending locations of reported structural variants. The measure is then defined as:

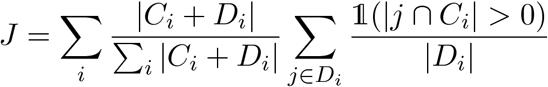

i.e., we weight the fraction of ground truth events that overlap with called events using the total mutation count over chromosomes. Notably, all of these measures have their range in the unit interval [0, 1] which allows for straightforward construction of more nuanced measures.

### 3.2 Statistical Test for Mutation Rate Variation

As a first proof-of-concept demonstration, we evaluate whether a given study design would be able to detect a difference in mutation rates between tumors for single nucleotide variation. A motivating hypothesis for these tests is the idea that cancerous and precancerous tissues should typically exhibit hypermutability phenotypes, that is elevated rates of particular kinds of variation that lead to genetic heterogeneity across cells. We would then wish to detect whether a specific study design would be powered to detect a hypothetical difference in mutation rate between two samples indicative of a hypermutability phenotype specific to one sample.

We assume that we have two independently sampled tissues, and we wish to determine whether the mutation rates of various mutation classes is different between these two tissues. To create a specific hypothetical scenario, we generated two sets of SNV data, one where the average rate was high and the other where it was comparatively low (denote the rates as *λ*_1_ *≈* 10^−8^, *λ*_2_ *≈* 10^−10^ mutations per nucleotide per generation). We specifically tested whether a 30x WGS screen on both tissues would be statistically powered to detect the mutation rate difference. All other parameters were kept equal at reasonable values (0 error rate, paired-end sequencing, 0 single cells, with 1 sample per tissue). We assume that the first tissue has mutation count generated as *M*_1_ ∼ *Poisson*(*λ*_1_*t*_1_) and the second tissue has mutation count generated as *M*_2_ ∼ *Poisson*(*λ*_2_*t*_2_). A hypothesis test that can be used to test for a rate difference is then represented by the following null and alternative hypothesis: 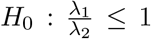 and 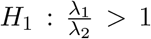. In our setting, we have an estimate of the mutation counts *M*_1_ and *M*_2_ from the output of our variant calling software, and we estimate *t*_1_ and *t*_2_ as biologically plausible tumor formation times as described below. Several statistics can be used to establish p-values for a two-sample Poisson rate test (c.f., (12)), but for the present example favor a conditional test based on the fact that the conditional distribution of *M*_1_ given *M*_1_ +*M*_2_ is binomial. Under the assumptions of the problem, this expression, which we denote *p*(*t*_1_, *t*_2_) to emphasize its dependence on each time, is:

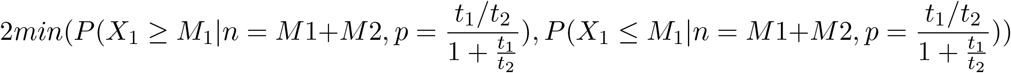

These values are both computed easily using binomial cdf computing packages.

In our empirical test case, 244 mutations were called in the higher SNV rate dataset and 2 were called in the lower SNV rate dataset. Times *t*_1_ and *t*_2_ are estimated, with uncertainty, as depths of somatic lineages which could plausibly lead to a tumor; we assign them random variables 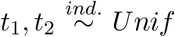 [1, 30] years. Current evidence suggests that the time from a normal cell to clinical cancer sequencing could be as high as many decades, though studies are in their nascence for somatic charting in healthy and precancerous tissues (15). To generate a p-value, we want the expectation of our p-value function with respect to the times, i.e.

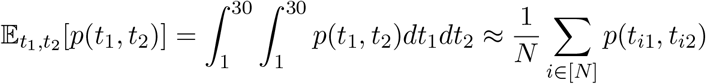

Since we lack an elementary closed form expression for *p*(*t*_1_, *t*_2_), we numerically approximate the integral via a bootstrap procedure. In our test case we took 500 random draws of times *t*_1_, *t*_2_ and computed an average p-value of 6.9 × 10^−7^ with a variance of 1.4 × 10^−10^. Therefore, we can conclude in this test case that the given study design should be powered to provide a strongly significant detection of the given rate variation.

### 3.3 Evaluating Study Parameter Choices

A more involved use of the simulator would be to test a range of study designs and identify those powered to detect a hypothesized effect. To provide a concrete example of such a study design question, we first evaluate how varying coverage would change our ability to call single nucleotide variants. To do this, we generated eight sets of data, each with an average SNV rate of approximately 10^−8^, and a lower rate of approximately 10^−10^ for deletion and inversion variants. The read lengths for the dataset were fixed at 125, the error rate at 0, zero single cells were generated, one tumor was generated with eight samples from that tumor, and paired-end whole genome sequencing was performed. All biological parameters were fixed. The depth of coverage parameter was sampled from the set {1, 2, 5, 10, 15, 25, 30} for each of the eight samples. The F1 score for variant calling as a function of depth of coverage is visualized in Figure 2, with the main takeaway being that a coverage below 15 has a significant negative effect on our ability to call SNVs, whereas more incremental gains can be seen above 15x coverage.

**Figure 2:**
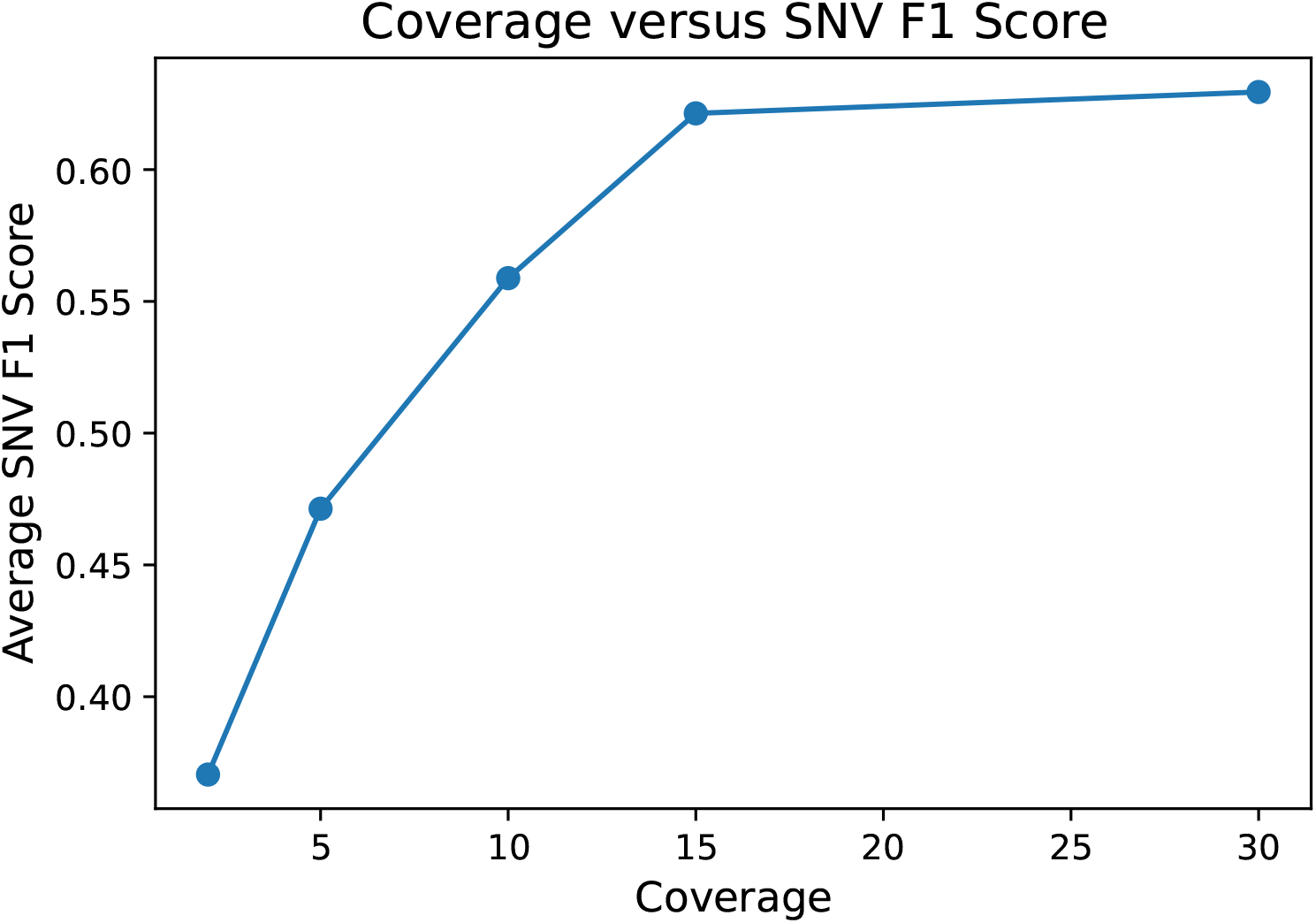
F1 score for SNV calling accuracy on simulated bulk sequencing data as a function of depth of coverage.

### 3.4 Optimizing a Study Design

The most involved intended use of the simulator is to optimize a study design to evaluate a particular hypothesis about somatic variability. Researchers may be interested in studying a particular attribute of a cancer to gain insight into the system. For instance, one might be interested in variability in chromothriptic events, while another could be interested in changes in clonal diversity. Our ability to answer each of these questions, or a combination of these questions, is dependent on study design choices made before data is gathered. For example, some study designs (e.g., if coverage is not sufficient) would not be sufficiently statistically powered to detect an accurate signal. At the same time, resource limitations will affect the feasibility of possible designs. The presumed goal is to design a study that is optimally powered to detect the hypothesized signal within available resource constraints. Here, we demonstrate the use of our simulator to answer such a study design question to find an approximately optimal study design for a particular hypothesis.

The prior premise can be framed as the following optimization problem:

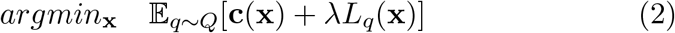

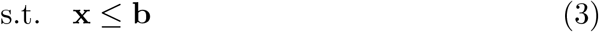

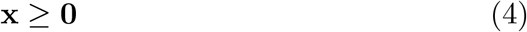

Here **x** is the study design vector defined in Equation (1), though in practice only a subset of the variables in **x** may be relevant. **c**(**x**) is a cost function for a particular study design and **b** is a maximal budget vector which constrains the amount of each resource in the study design we can use. Constraint (4) enforces non-negativity of each of the study design variables. We also define an unknown loss function *L*_*q*_(**x**), which describes the error with which the study design answers the question we pose. *L*_*q*_(**x**) is generally our focus, and is a flexible function dependent on the questions we want to answer. If we were concerned with phylogenetic structure and variant calling accuracy, our loss function could be some combination of tree inference and variant calling precision. If we were concerned with mutational rate differences across tissues, we could use some loss function derived from a hypothesis test. The *λ* tuning parameter determines the amount with which we prioritize this loss function. The subscript *q* and the expectation term over the distribution *Q* are used to emphasize that instances of our simulator are evaluated on a biological parameter vector *q* drawn from a stochastic high-dimensional biological parameter distribution *Q*. The number of clones, clonal frequency in each sample, mutation rates/types, and mutation locations are all part of this distribution *Q*, and each simulator run draws some particular value *q*_*i*_ ∼ *Q* which we evaluate with the loss function. Since we lack a closed form expression for the distribution *Q*, the expectation can be approximated by Monte-Carlo methods, where a singular study design, **x** would be evaluated on multiple instances *q*_*i*_, and the result averaged, i.e. 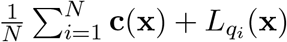

To provide a tractable test case for optimal study design, we assume that our study design variable **x** can vary only in numerical read length, coverage, and error rate, and a binary decision of whole genome versus whole exome sequencing. We assume the study is designed to recover inversions, deletions, and SNVs in a sample, so we define a scoring function intended to provide a balanced measure of performance at these tasks:

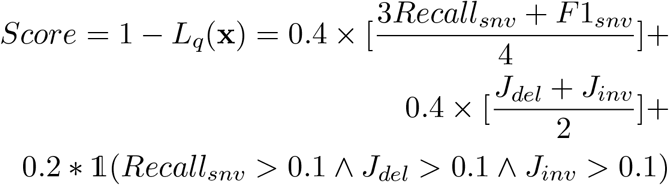

This scoring function gives a greater priority to identifying correct variants by over-weighting recall; it also provides a “bonus” for study designs able to detect all three types of variation at a significant rate. We fix our callers as `strelka` and delly, fix most biological parameters, and fix our number of samples at 1 (i.e. no multi or merging sampling). We assume that all study designs have fixed cost **c**(**x**) = 0 and constrain our study design as follows: *rl* ∈ [75, 2000]; *c* ∈ [2, 30], *e* ∈ [0, 0.001], 1(*Paired*) ∈ {*True*}, 1(*Genome*) ∈ {*True, False*}, *n* ∈ {0}. We replicated tumors independently with fixed biological parameters, fixing *k* = 5, *N*_*e*_ = 8*e*8, *α* = 10, and sampling SNVs, inversions, and deletions with rates in the range [10^−14^, 10^−8^]. For evaluation, approximately 75 study design vectors **x** were generated and evaluated from around 15 tumors. The score function was averaged across all tumors with respect to study design to generate a final score for each study design.

Our top 3 choices for the constrained optimal study design were: (2000, 25, 0.0, *True*), (2000, 30, 0.001, *True*), (2000, 10, 0.0, *True*), where we represent the study design as the vector (*rl, c, e*, 1(*Genome*)). The best and worst four non-zero score study designs are reported in Table 4. As expected, the best study designs were mostly whole genome sequenced, which allowed the recovery of more variants across noncoding regions of the genome. We might expect that resource constraints or a scoring function that penalized for the higher resource usage of whole-exome sequencing could reach different conclusions. Similarly, high coverage did seem to boost the power of the design, but there did not appear to be a large difference between 10x and 30x coverage. Extremely low coverage, however, did result in poor scores. Increasing the read length boosted our ability to detect structural variants in samples, in both exome and genome sequenced samples. The error rate parameter did impact the rate of false calls and the ability to detect SNVs, but the overall scores were not heavily affected by the error rate. This was somewhat by design, as we mainly prioritized the sensitivity to detect variants in a sample as opposed to accounting for false positives. As expected, we saw three orders of magnitude more false SNV calls in 0.001 error rate samples as compared to lower error values. However, with respect to structural variation, the larger error rate samples did not appear to do significantly worse. This meant that long-read high error-rate designs did well with respect to our scoring measure. The worst study designs were often exome-sequenced, had poor coverage, or completely failed to recover a particular variant type due to noise. Figure 3 visualizes tradeoffs between score and the model design parameters in our exploration of the search space.

**Table 4:** Top Four Scoring and Bottom Four(non-zero) Scoring Study Designs

| Read Length | Coverage | Error Rate | Exome Sequenced | *Score* |
| --- | --- | --- | --- | --- |
| 2000 | 25 | 0.0 | False | 0.587395 |
| 2000 | 30 | 0.001 | False | 0.496574 |
| 2000 | 10 | 0.0 | False | 0.391434 |
| 150 | 25 | 0.0 | False | 0.283998 |
| Read Length | Coverage | Error Rate | Exome Sequenced | *Score* |
| 500 | 2 | 0.0001 | True | 0.001818 |
| 150 | 30 | 0.001 | True | 0.005085 |
| 75 | 10 | 0.00001 | True | 0.011801 |
| 2000 | 2 | 0.00001 | True | 0.018933 |

**Figure 3:**
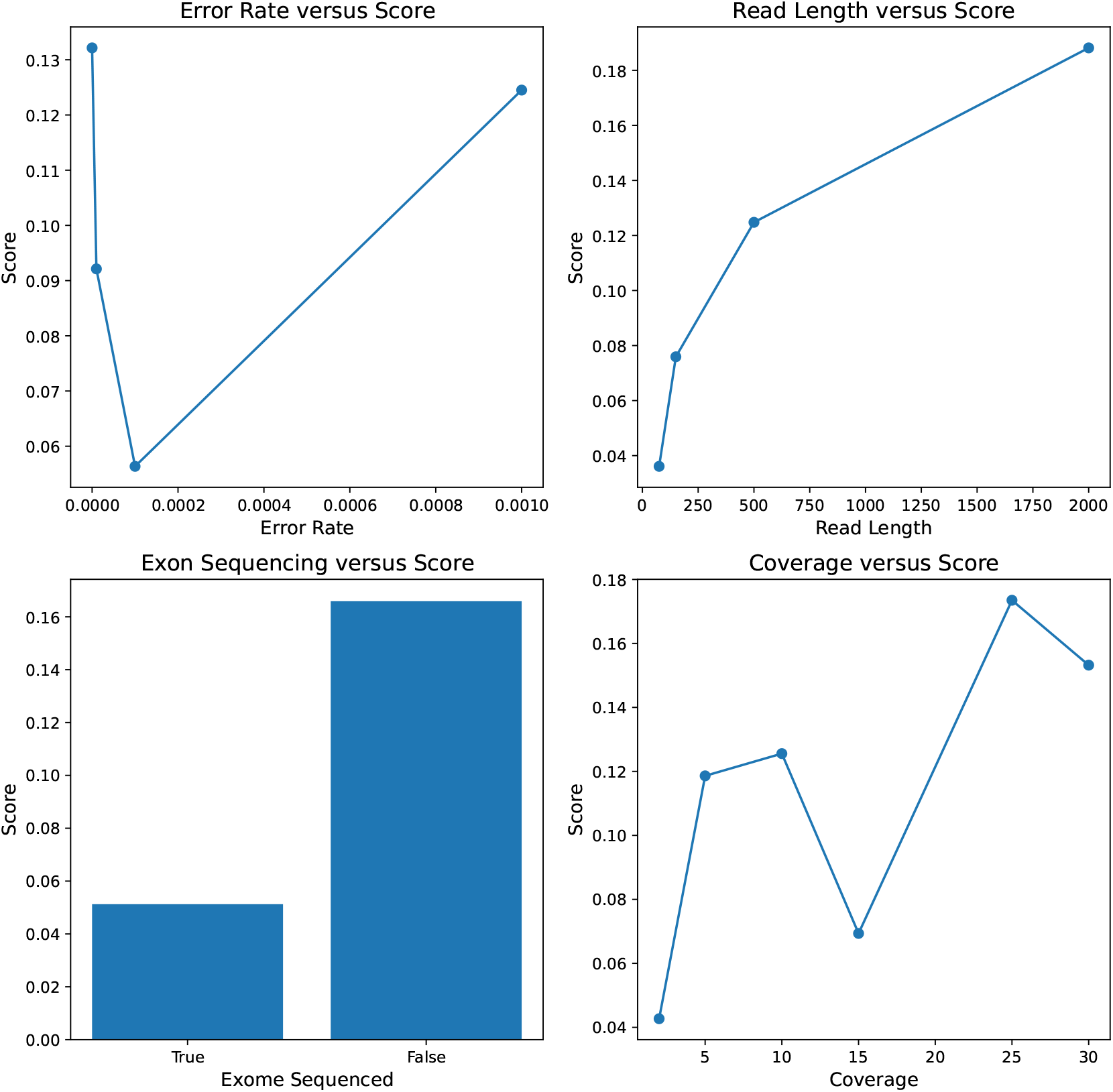
Visualizations of various parameters and their impact on the efficacy of a study design. Read length showed the strongest positive correlation with study design score, and similarly whole genome sequenced data had a higher score than exome sequenced data. Higher coverage had a generally positive, but not entirely consistent, effect on the study design score. Higher error rates also generally produced lower scoring study designs, but high error rates did not preclude a study from having a high score.

Aside from finding useful study design vectors and study design parameters, our simulations yielded a number of tangentially interesting results. One finding that may have implications for future cancer informatics development was the effect of hypermutability and rearrangements on our ability to recall variants. High frequency structural events (e.g. deletions and inversions) introduced a substantial amount of noise in our SNV caller report. While this may be expected due to the sensitivity of the `Strelka` caller, it also may have ramifications in the clinical use case. The simulator may also be over-reporting these false calls by placing a uniformly high quality score on all bases in the simulation process since presumably bases with errors may have lower quality scores which may then be accounted for by the variant software. As expected, the task of calling layered structural variation at various frequencies was more challenging than that of single nucleotide variation. Predefined coordinates with respect to a linear genome may not be the most effective way to classify these events; other abstractions that consider temporal orderings of various rearrangement events in flexible genome collections may provide further insight into the nature of mutability. For example, Zeira et al. (48) have made progress in this direction of identifying sequences of rearrangement events that generate tumor genomes. Regardless of analysis type, the patterns that large scale hypermutability, high error, and imbalanced clonal samples can induce on sequencing reads should likely be considered when developing the next generation of cancer informatics tools. A final unrelated note is that a brief examination of variant reports reinforces common variant calling practices with respect to ensembling, i.e. merging the results of calls on independent samples increased the recall of variants present in the tumor. However, care must be taken as a number of false calls were introduced, especially with higher error rate sequencing.

### 3.5 Comparison to Real Cancer Reads

We finally sought to examine the ways in which our simulated reads were similar to and different from those produced by a real sequencing study. For this evaluation, we used an Ion Torrent targeted cancer sequencing panel in a colorectal cancer patient, found in the sequencing read archive(SRA Access Key: SRX9731615). This particular case used single end reads in the read-length range 50-300 base pairs. For a comparison case, we generated 150 base pair exome-sequenced reads. The GC content distributions as well as the GC percentage of the reads in the real reads seemed to closely mirror that in our simulated reads. Namely the real reads had a GC content of 46 percent whereas ours had a GC content of 44 percent. This was somewhat surprising as one might expect the GC content of real sequencing reads to differ somewhat from our uniform fragmenting procedure. However, this observation seems to make sense in light of the fact that GC bias seems more pronounced for Illumina sequencing platforms (3). The read length distribution for the Ion Torrent reads were more spread out than in our simulated data, having a range of [25, 354], whereas ours ranged in [0, 150] by design. In our sequencing case, we did two forms of truncation on the fragments and reads, not generating fragments that were too small (except at the end of chromosomes), and also only writing sequences of the prescribed length on that fragment. The distributions over read lengths can be adjusted in our simulation to allow for broader, and uncapped, ranges. The final noteworthy difference was with respect to quality scores. In our simulator we placed a uniform quality score on all bases. While this is a naive assumption it did not prove to be extremely different than that of the real sequencing case. In particular, the real sequencing case had fairly uniform sequencing quality scores for bases until the reads became extremely long, after which point the quality dropped.

While the simulated read files do not exactly mirror the distributions found in current sequencing technology platforms, we are mainly concerned with invariance with respect to sequencing attributes. That is, a shift in signal accuracy caused by simulated sequencing parameters (say, read length), should produce a similar shift in accuracy in real sequencing technologies under that same shift in sequencing parameters. In a loose sense, our simulated reads can be viewed as a limiting case of current sequencing platforms, which does not over-represent parts of the genome and has a more defined distribution of read lengths. This is not necessarily an issue for the purpose of this simulator, as it is conceivable that future technologies/sequencing protocols could be developed that do not possess the same read length distributions, read overrepresentations, or sequencing quality scores. Additionally, these distributions vary depending on the sequencer and random stochasticity – they could, however, be plausibly integrated into the simulator in future iterations.

## 4 Discussion and Conclusion

In this paper, we introduced a new simulation toolkit to generate sequencing reads from somatic variation processes under a wide range of biological and technological parameters. We used a bottom-up approach, encoding various aspects of somatic evolution and sequencing with user customizable probability densities. We demonstrated the utility of our simulator for several hypothetical questions in evaluating and optimizing study designs for profiling somatic variability in cell lineages.

A downside of our approach is that in the service of modeling general classes of technologies, we may not encode some unique properties of specific sequencing platforms. Details such as distributions over sequencing quality and exon error tolerance are somewhat crudely approximated by our simulator and might need to be customized to specific current platforms in future work. In the exon sequencing case, we try to find subsets of reads that may match exon sequences well, but this is done in an inexact way due to computational considerations. Another approximation in our experimental tests was the de facto alignment to a linear reference genome as the first part of our experiments. In the case of highly rearranged cancer genomes, alignments may not provide high quality insights into the original mutation sources. Alternatives such as graph-based alignments or reference-free sketching ideas that could be explored in the future. In the case of ultra-long reads, it may be computationally feasible to assemble the genome, raising further questions not explored here such as the lengths at which assembly becomes feasible.

The simulator might also be extended in various ways in future work. While DNA sequencing has come to be the standard lens by which researchers view the cancer evolutionary system, a growing body of work on epigenetic theory demonstrates that some neoplasms may use epigenetic modifications to generate a selective advantage (16). Incorporating various forms of epigenetic modifications – 3D genome alterations, methylation, etc.– and the technological methods used to probe these changes could be a valuable addition to our simulator. As our knowledge of the mechanisms of somatic evolution and mutagenesis change, modifications could be made to our simulation system to incorporate these novel patterns. For instance, progress in understanding complex patterns of structural variations and the mechanisms by which they arise in somatic cells is still in its infancy, with new patterns of structural variation continuing to be discovered (13). It may be possible for future iterations of the simulator to encode arbitrary rearrangements in the genome rather than those that come in defined arrangements. Additionally, novel mutational signatures are being discovered with a wide variety of endogenous and exogenous causes. Novel mutational signatures and alternative forms of structural variation could be readily incorporated into our current framework by modifying the distributions over genome lengths and frequencies of mutation. Evolutionary modeling is another potential area of improvement. We utilized a neutral coalescent model to represent the evolutionary process stemming from a single cell, and a Dirichlet process to model the clonal frequencies in each sample. There is room here to incorporate various selection pressures, clonal dynamics, drift, and bottleneck effects with greater knowledge of how these processes act in the cancerous setting.

The primary goal of this simulator is to allow thorough exploration and optimization of spaces of study design decisions and evaluate their impacts on our power to detect significant patterns of somatic evolution. We are particularly interested in our ability to reconstruct evolutionary lineages, find their characteristic mutational signatures and rates, and detect patterns of structural variation. An important task going forward is to provide user-friendly software for study-design inquiries. This software would allow a user to input properties they wish to detect in a cancer sample along with cost settings; the software would then return sets of study design parameters which allow for their detection under minimal cost. Ideas from Bayesian hyper-parameter optimization will likely prove useful in our optimization goals since each iteration of our output function is expensive to obtain. Ideally, we wish for a symbiotic loop between sequencing technology development and simulation study-design optimization. That is, simulations could produce realistic sets of data of a neoplastic process, optimization techniques could then produce feasible sets of technological parameters with which details of this process are revealed, and finally sequencing technological development could then be targeted towards parameter sets that provide maximal amounts of information.

## 5 Acknowledgements

We thank the AWS ML Research Awards program for providing cloud computing support for this work.

## Data Availability

Simulated data created for this study are provided with the source code at https://github.com/CMUSchwartzLab/MosaicSim

## A Appendix

### Algorithm 1

Pseudocode for Read Generation for Whole Genome Sequencing

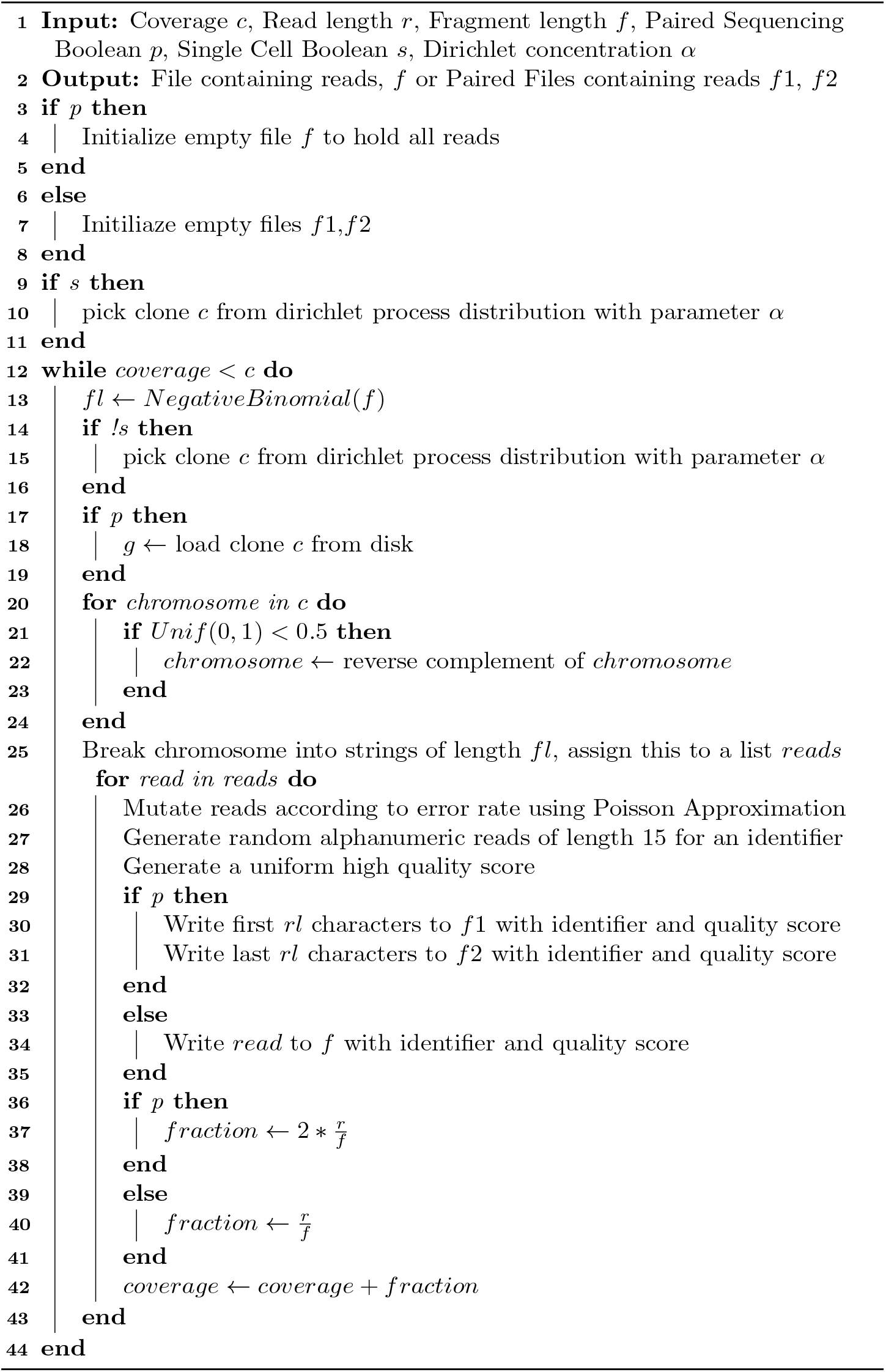

### Algorithm 2

Pseudocode for Read Generation for Whole Exome/Targeted Sequencing

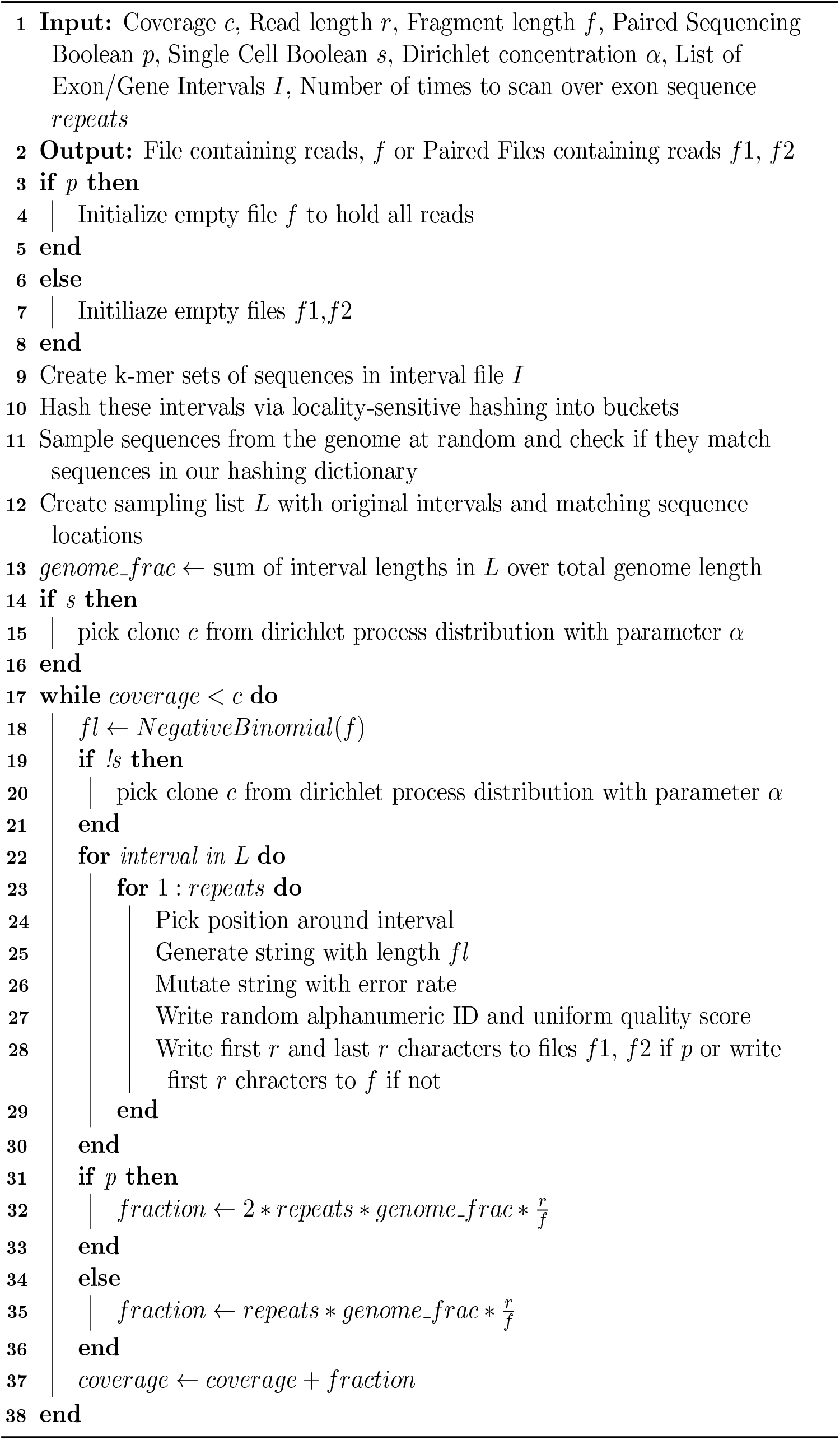

### Algorithm 3

Generate Mutational Events

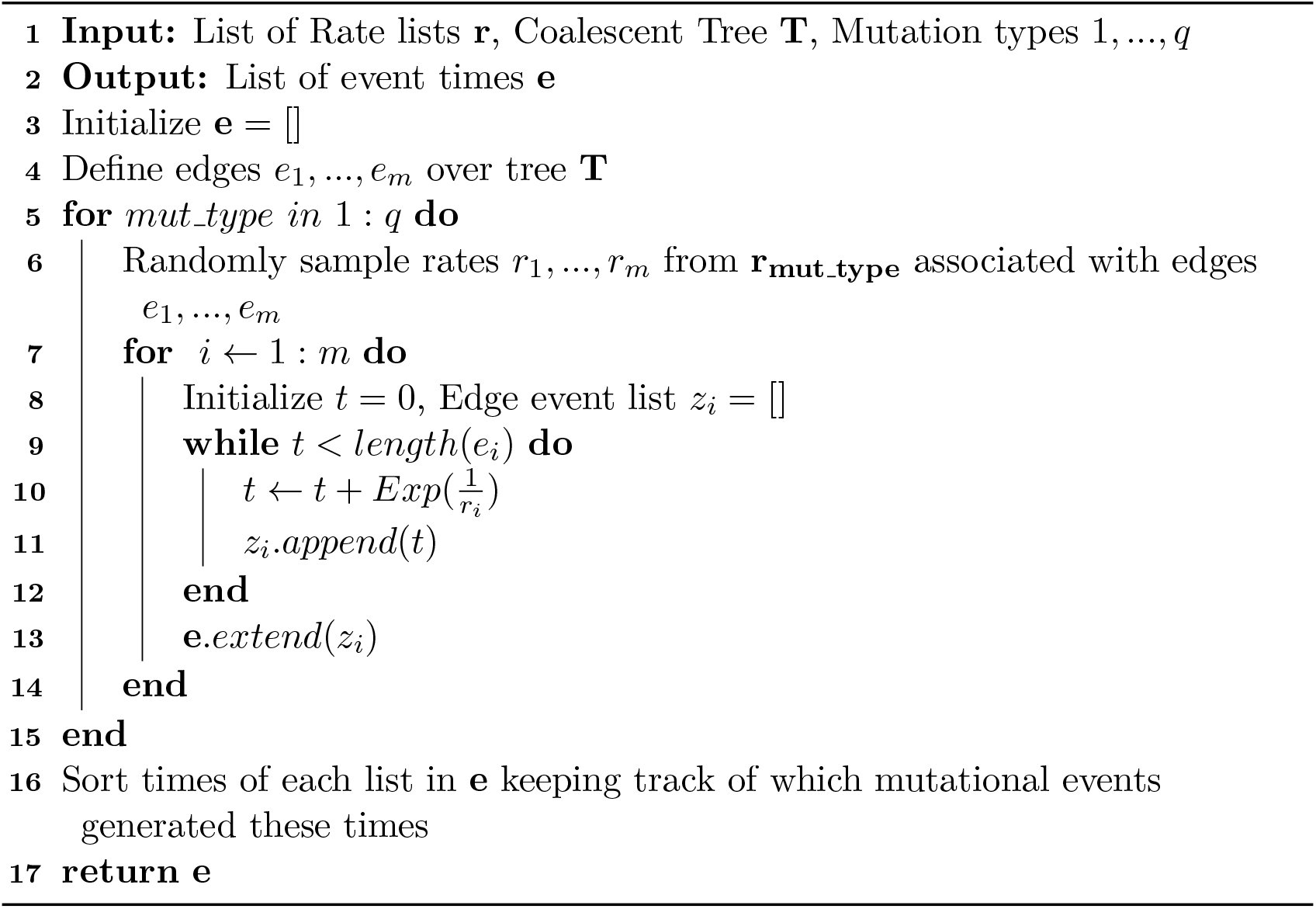

### Algorithm 4

Generate Clonal Genomes

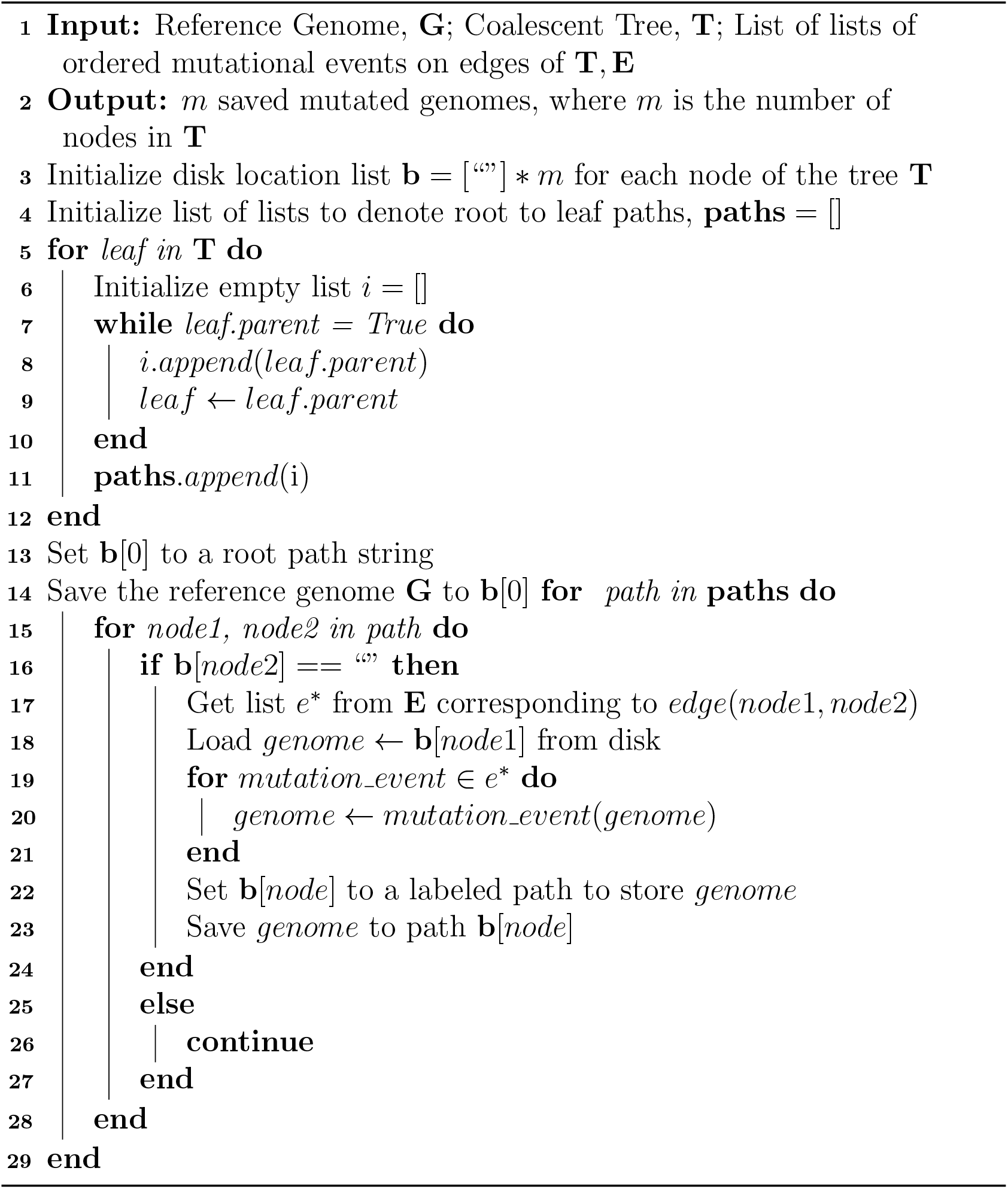

### Algorithm 5

Single Nucleotide Polymorphism Implementation

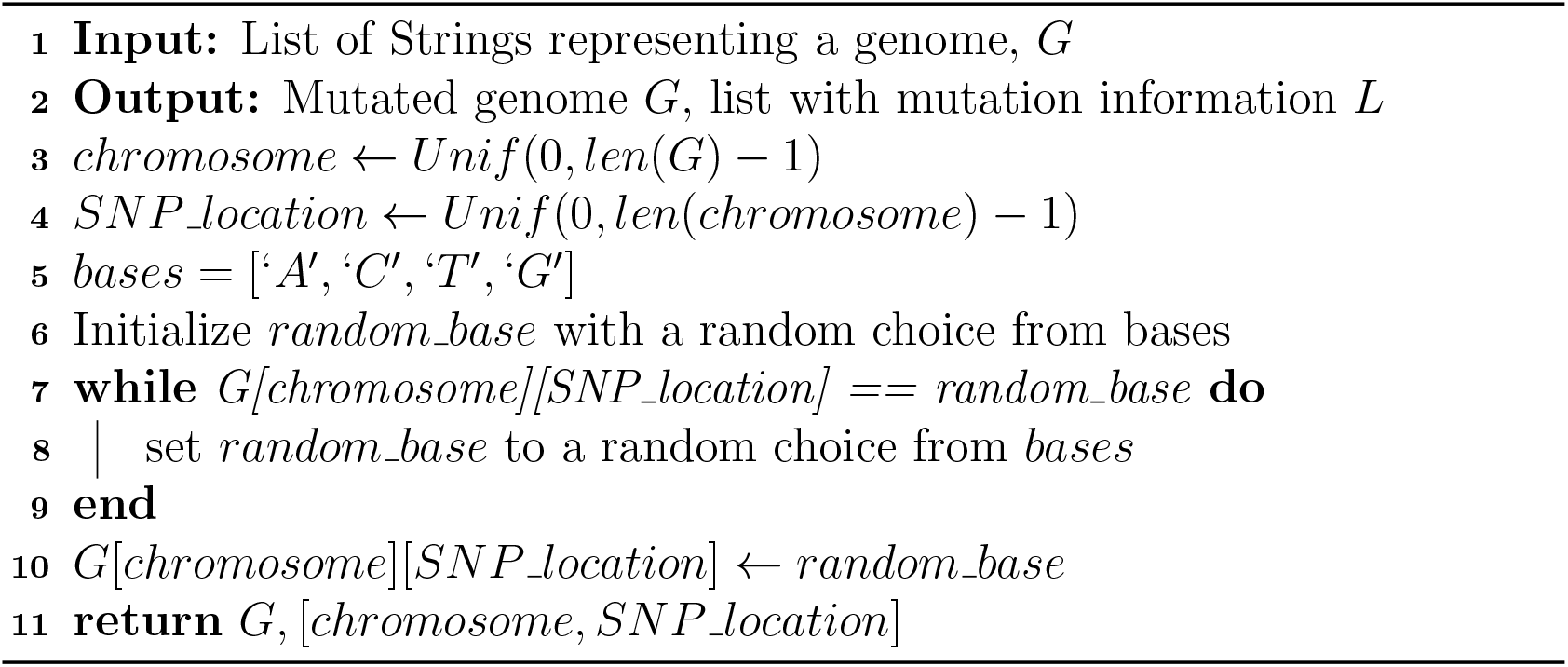

### Algorithm 6

Copy Number Variation Implementation

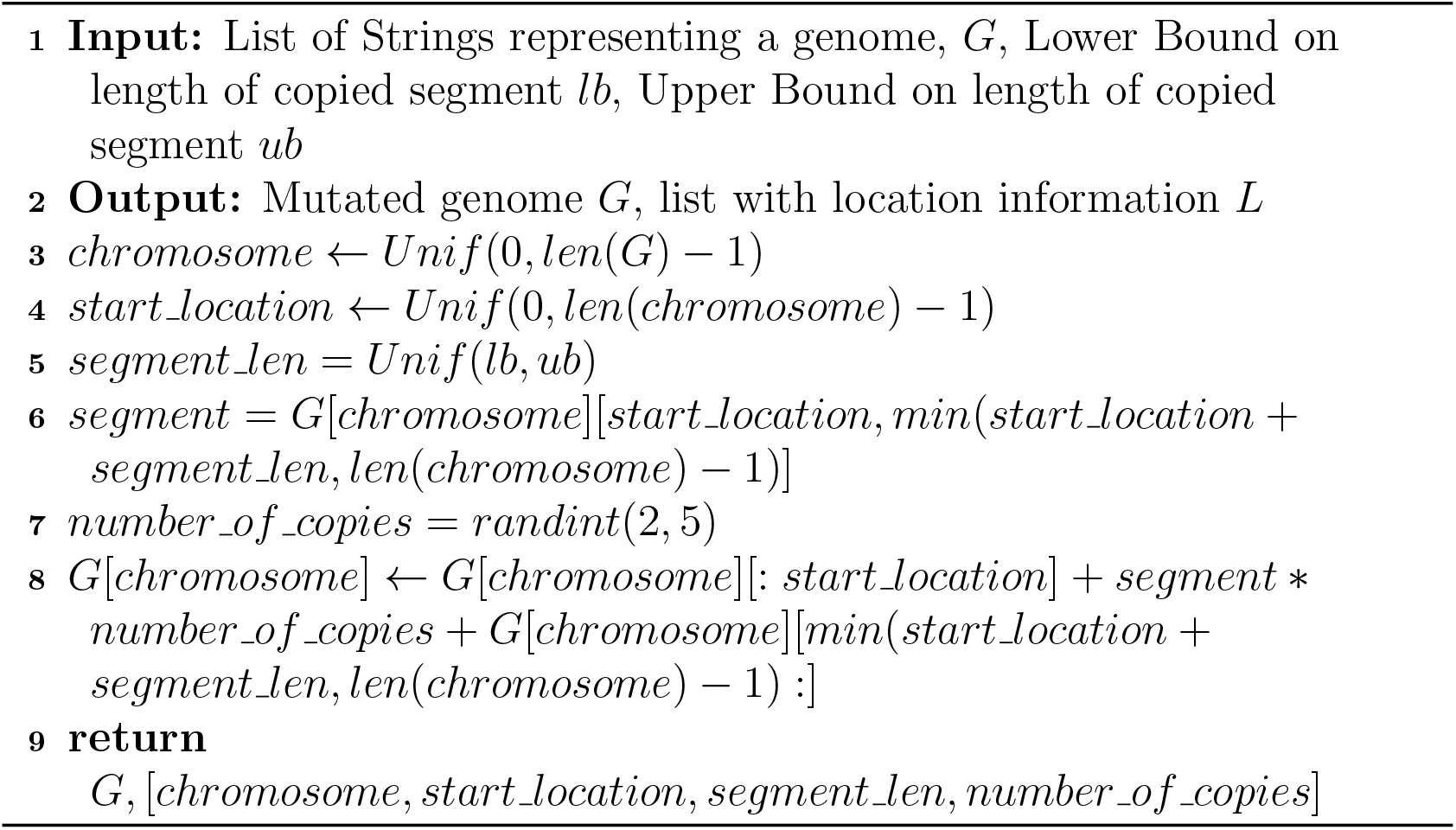

### Algorithm 7

Deletion Implementation

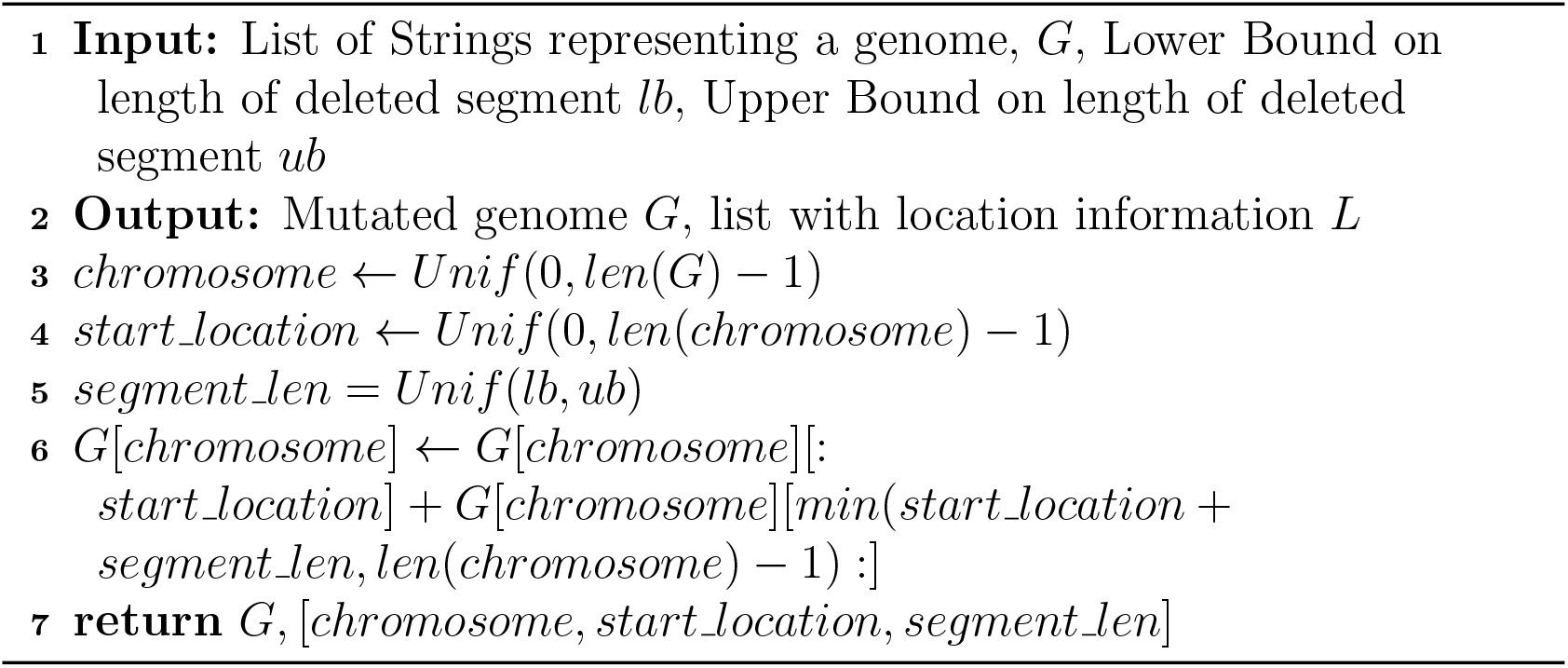

### Algorithm 8

Inversion Implementation

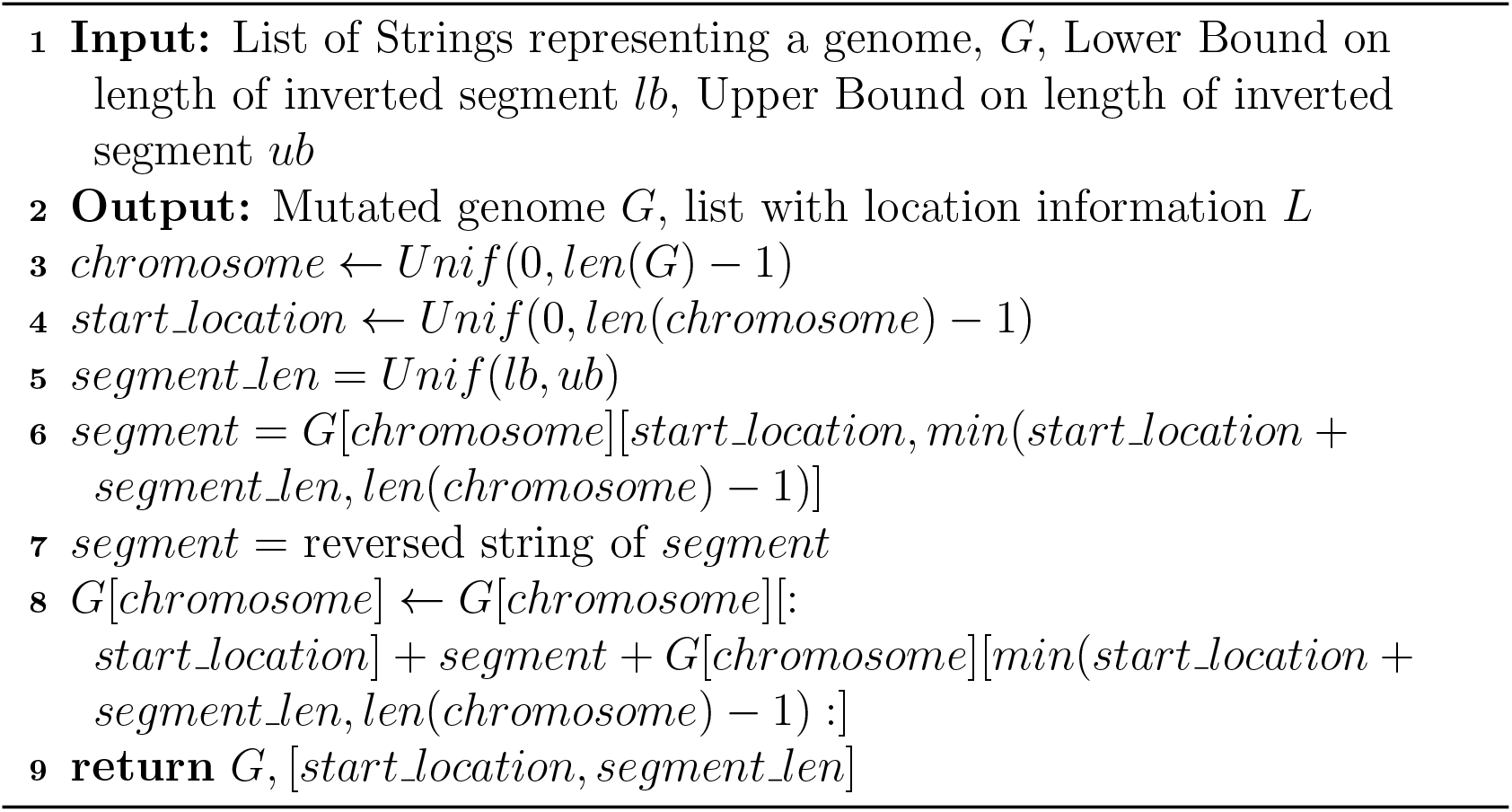

### Algorithm 9

Aneuploidy Implementation

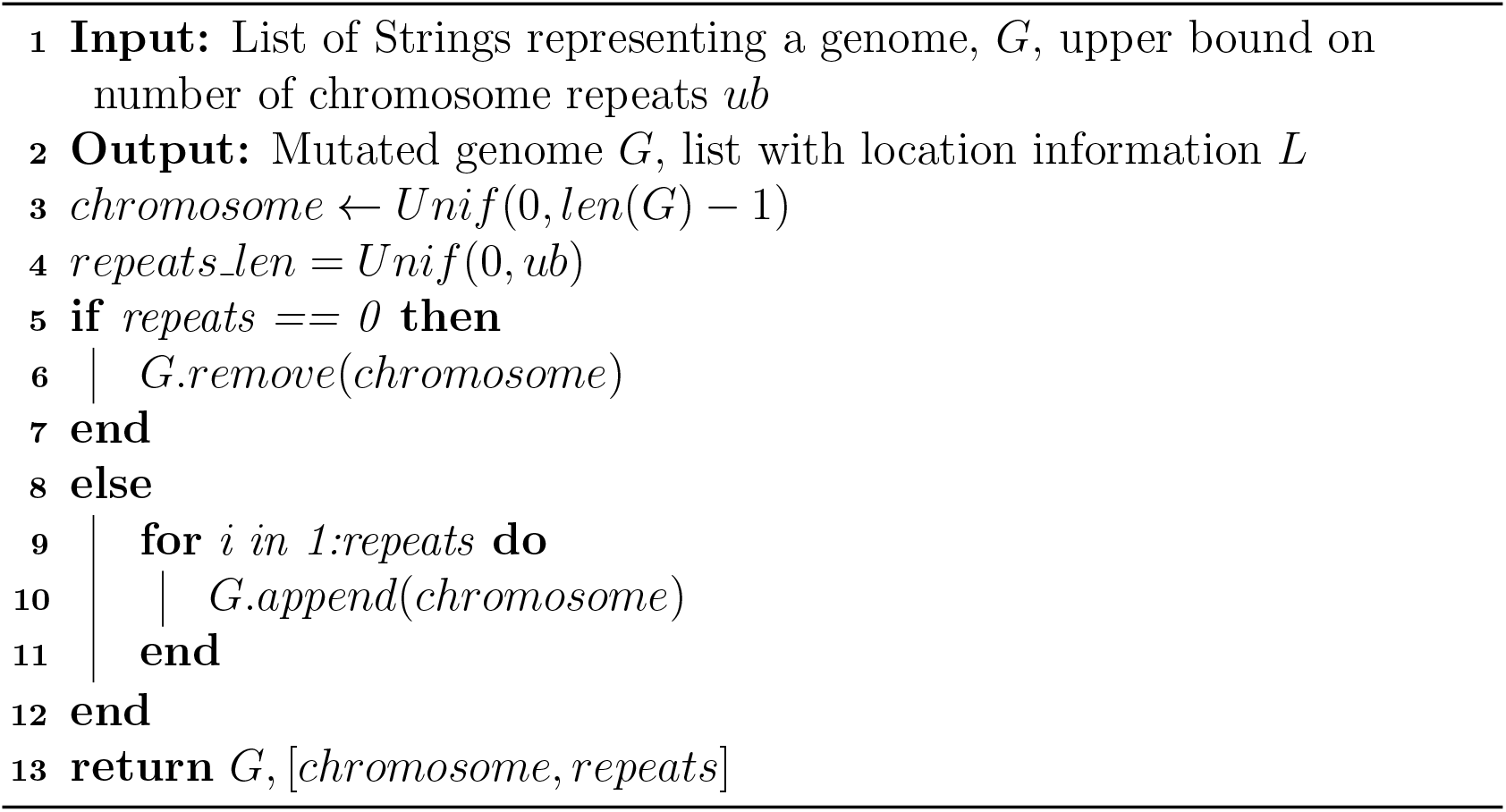

### Algorithm 10

Insertion Implementation

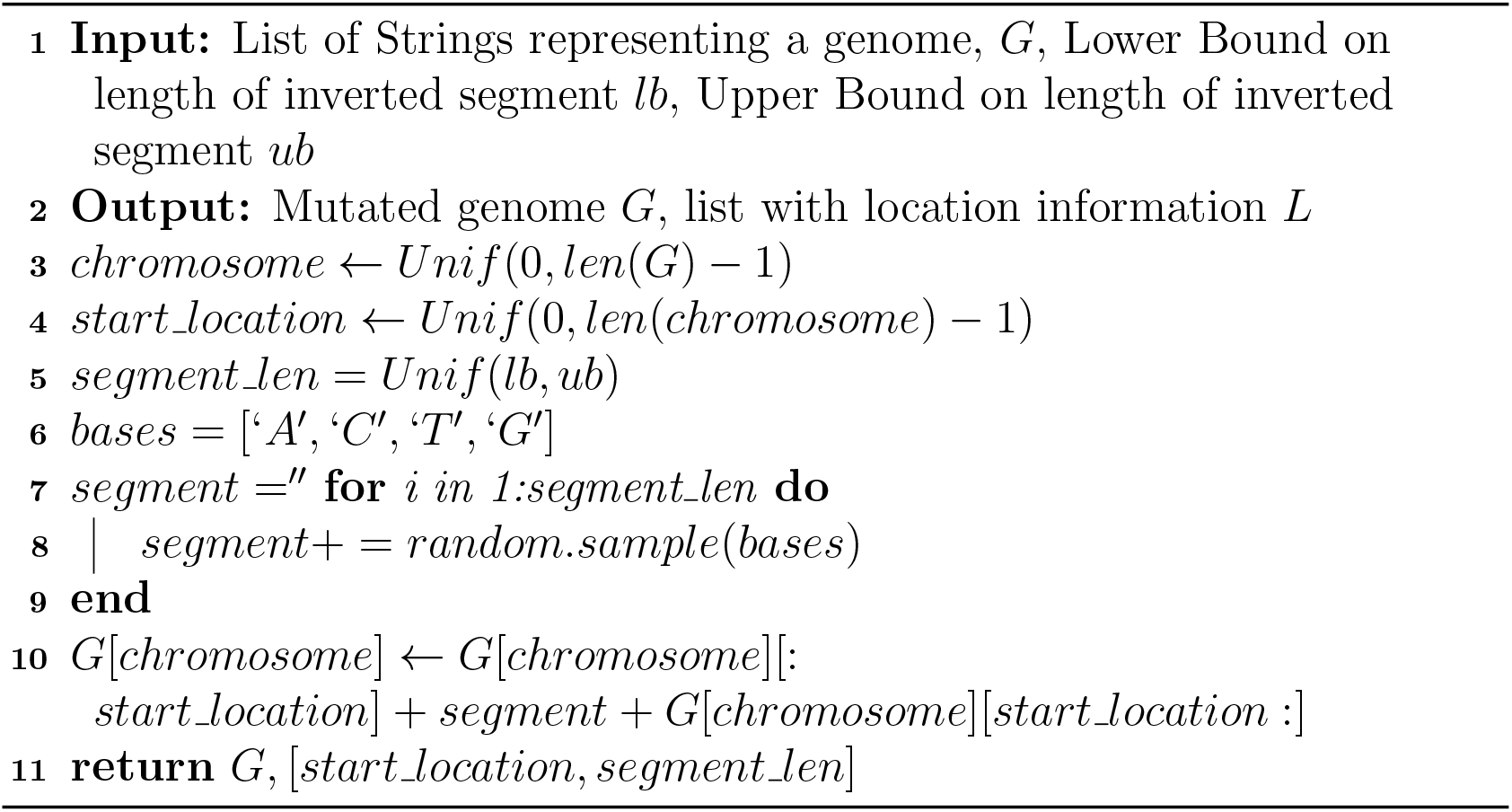

### Algorithm 11

Chromothripsis Implementation

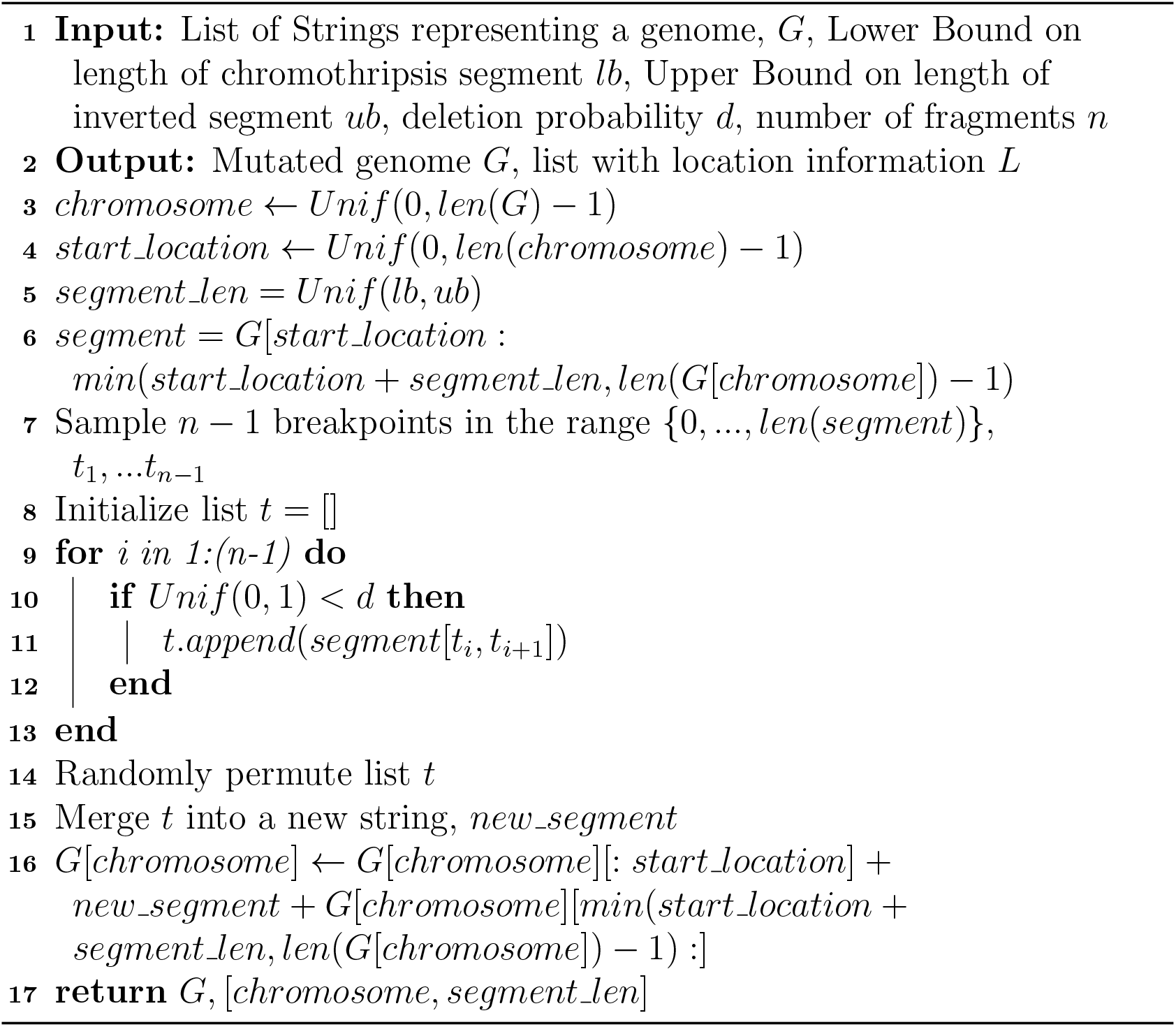

### Algorithm 12

Kataegis Implementation

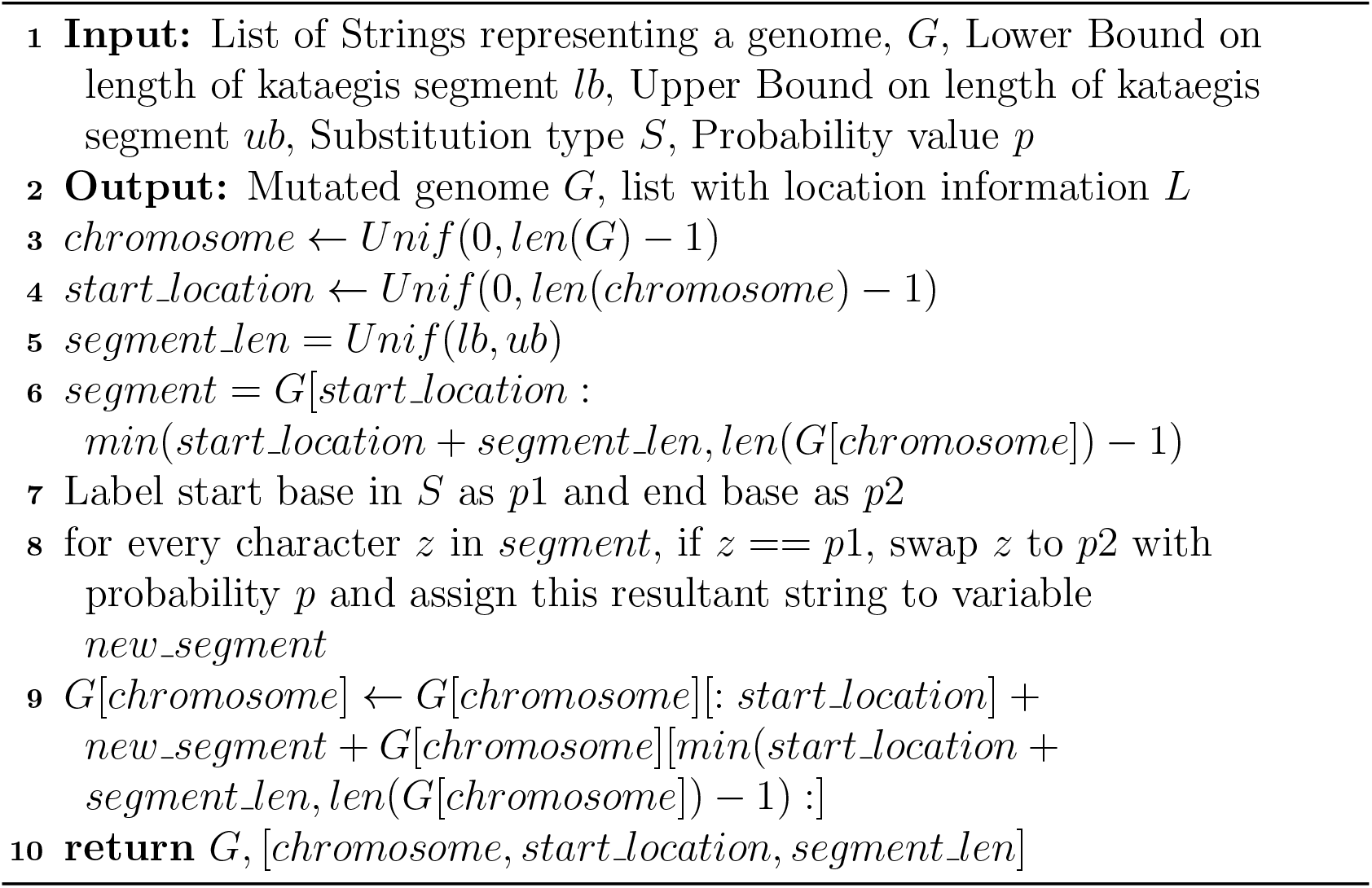

### Algorithm 13

Breakage Fusion Bridge Implementation: TODO

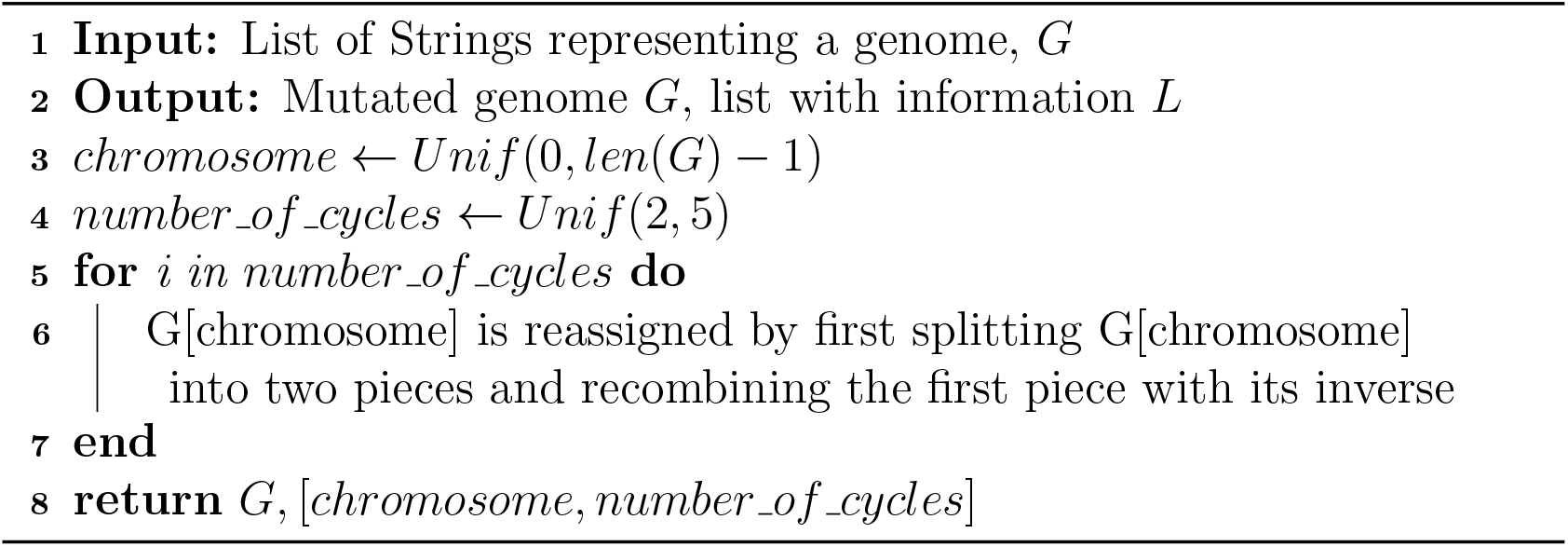

### Algorithm 14

Chromoplexy Implementation

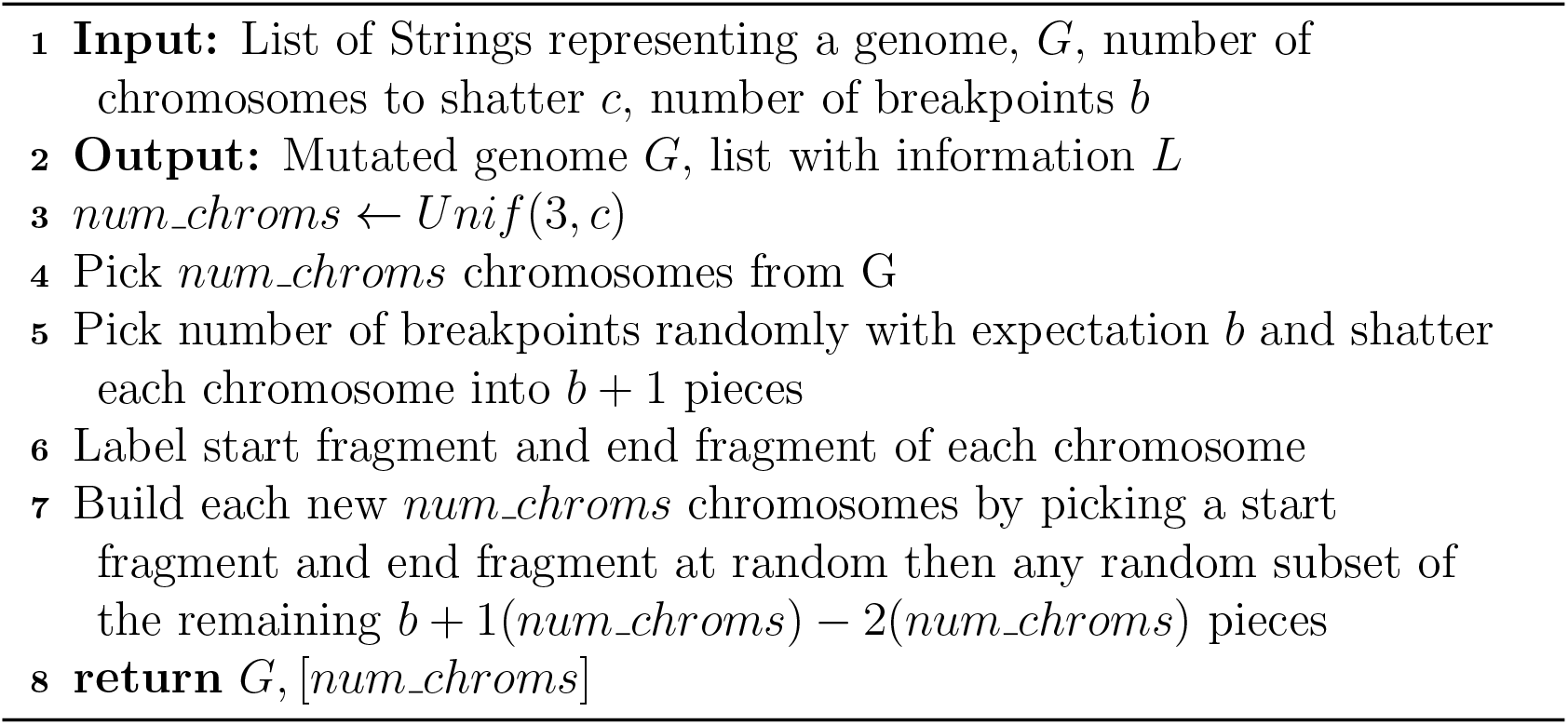

## References

[1] Abascal, F., Harvey, L.M., Mitchell, E., Lawson, A.R., Lensing, S.V., Ellis, P., Russell, A.J., Alcantara, R.E., Baez-Ortega, A., Wang, Y., et al.: Somatic mutation landscapes at single-molecule resolution. Nature 593(7859), 405–410 (2021)

[2] Alexandrov, L.B., Kim, J., Haradhvala, N.J., Huang, M.N., Ng, A.W.T., Wu, Y., Boot, A., Covington, K.R., Gordenin, D.A., Bergstrom, E.N., et al.: The repertoire of mutational signatures in human cancer. Nature 578(7793), 94–101 (2020)

[3] Benjamini, Y., Speed, T.P.: Summarizing and correcting the gc content bias in high-throughput sequencing. Nucleic acids research 40(10), e72–e72 (2012)

[4] Colom, B., Herms, A., Hall, M., Dentro, S., King, C., Sood, R., Alcolea, M., Piedrafita, G., Fernandez-Antoran, D., Ong, S., et al.: Mutant clones in normal epithelium outcompete and eliminate emerging tumours. Nature pp. 1–5 (2021)

[5] Coorens, T.H., Moore, L., Robinson, P.S., Sanghvi, R., Christopher, J., Hewinson, J., Przybilla, M.J., Lawson, A.R., Spencer Chapman, M., Cagan, A., et al.: Extensive phylogenies of human development inferred from somatic mutations. Nature 597(7876), 387–392 (2021)

[6] Dentro, S.C., Leshchiner, I., Haase, K., Tarabichi, M., Wintersinger, J., Deshwar, A.G., Yu, K., Rubanova, Y., Macintyre, G., Demeulemeester, J., et al.: Characterizing genetic intra-tumor heterogeneity across 2,658 human cancer genomes. Cell 184(8), 2239–2254 (2021)

[7] Dou, Y., Gold, H.D., Luquette, L.J., Park, P.J.: Detecting somatic mutations in normal cells. Trends in Genetics 34(7), 545–557 (2018). https://doi.org/https://doi.org/10.1016/j.tig.2018.04.003, https://www.sciencedirect.com/science/article/pii/S0168952518300738

[8] Ellis, P., Moore, L., Sanders, M.A., Butler, T.M., Brunner, S.F., Lee-Six, H., Osborne, R., Farr, B., Coorens, T.H., Lawson, A.R., et al.: Reliable detection of somatic mutations in solid tissues by laser-capture microdissection and low-input dna sequencing. Nature Protocols 16(2), 841–871 (2021)

[9] Escalona, M., Rocha, S., Posada, D.: A comparison of tools for the simulation of genomic next-generation sequencing data. Nature Reviews Genetics 17(8), 459–469 (2016)

[10] Ewing, A.D., Houlahan, K.E., Hu, Y., Ellrott, K., Caloian, C., Yamaguchi, T.N., Bare, J.C., P’ng, C., Waggott, D., Sabelnykova, V.Y., et al.: Combining tumor genome simulation with crowdsourcing to benchmark somatic single-nucleotide-variant detection. Nature methods 12(7), 623–630 (2015)

[11] García-Nieto, P.E., Morrison, A.J., Fraser, H.B.: The somatic mutation landscape of the human body. Genome biology 20(1), 1–20 (2019)

[12] Gu, K., Ng, H.K.T., Tang, M.L., Schucany, W.R.: Testing the ratio of two poisson rates. Biometrical Journal: Journal of Mathematical Methods in Biosciences 50(2), 283–298 (2008)

[13] Hadi, K., Yao, X., Behr, J.M., Deshpande, A., Xanthopoulakis, C., Tian, H., Kudman, S., Rosiene, J., Darmofal, M., DeRose, J., et al.: Distinct classes of complex structural variation uncovered across thousands of cancer genome graphs. Cell 183(1), 197–210 (2020)

[14] Helleday, T., Eshtad, S., Nik-Zainal, S.: Mechanisms underlying mutational signatures in human cancers. Nature Reviews Genetics 15(9), 585–598 (2014)

[15] Jolly, C., Van Loo, P.: Timing somatic events in the evolution of cancer. Genome biology 19(1), 1–9 (2018)

[16] Kanwal, R., Gupta, S.: Epigenetic modifications in cancer. Clinical genetics 81(4), 303–311 (2012)

[17] Kelleher, J., Etheridge, A.M., McVean, G.: Efficient coalescent simulation and genealogical analysis for large sample sizes. PLoS computational biology 12(5), e1004842 (2016)

[18] Killcoyne, S., Yusuf, A., Fitzgerald, R.C.: Genomic instability signals offer diagnostic possibility in early cancer detection. Trends in Genetics (2021)

[19] Kim, S., Scheffler, K., Halpern, A.L., Bekritsky, M.A., Noh, E., Källberg, M., Chen, X., Kim, Y., Beyter, D., Krusche, P., et al.: Strelka2: fast and accurate calling of germline and somatic variants. Nature methods 15(8), 591–594 (2018)

[20] Koboldt, D.C.: Best practices for variant calling in clinical sequencing. Genome Medicine 12(1), 1–13 (2020)

[21] Koh, G., Degasperi, A., Zou, X., Momen, S., Nik-Zainal, S.: Mutational signatures: emerging concepts, caveats and clinical applications. Nature Reviews Cancer 21(10), 619–637 (2021)

[22] Langmead, B., Salzberg, S.L.: Fast gapped-read alignment with bowtie 2. Nature methods 9(4), 357–359 (2012)

[23] Lee, A.Y., Ewing, A.D., Ellrott, K., Hu, Y., Houlahan, K.E., Bare, J.C., Espiritu, S.M.G., Huang, V., Dang, K., Chong, Z., et al.: Combining accurate tumor genome simulation with crowdsourcing to benchmark somatic structural variant detection. Genome biology 19(1), 1–15 (2018)

[24] Li, H.: Aligning sequence reads, clone sequences and assembly contigs with bwa-mem. arXiv preprint 1303.3997 (2013)

[25] Li, H.: Minimap2: pairwise alignment for nucleotide sequences. Bioinformatics 34(18), 3094–3100 (2018)

[26] Li, Y., Roberts, N.D., Wala, J.A., Shapira, O., Schumacher, S.E., Kumar, K., Khurana, E., Waszak, S., Korbel, J.O., Haber, J.E., et al.: Patterns of somatic structural variation in human cancer genomes. Nature 578(7793), 112–121 (2020)

[27] Loeb, L.A.: A mutator phenotype in cancer. Cancer research 61(8), 3230–3239 (2001)

[28] Loeb, L.A., Bielas, J.H., Beckman, R.A.: Cancers exhibit a mutator phenotype: clinical implications. Cancer research 68(10), 3551–3557 (2008)

[29] Luzzatto, L., Pandolfi, P.P.: Causality and chance in the development of cancer. N Engl J Med 373(1), 84–88 (2015)

[30] Mallory, X.F., Edrisi, M., Navin, N., Nakhleh, L.: Assessing the performance of methods for copy number aberration detection from single-cell dna sequencing data. PLoS computational biology 16(7), e1008012 (2020)

[31] McTavish, E.J., Pettengill, J., Davis, S., Rand, H., Strain, E., Allard, M., Timme, R.E.: Treetoreads-a pipeline for simulating raw reads from phylogenies. BMC bioinformatics 18(1), 1–7 (2017)

[32] Metzker, M.L.: Sequencing technologies—the next generation. Nature reviews genetics 11(1), 31–46 (2010)

[33] Nicol, P.B., Barabási, D.L., Asiaee, A., Coombes, K.R.: Sith: an r package for visualizing and analyzing a spatial model of intratumor heterogeneity. bioRxiv (2020)

[34] Nordborg, M.: Coalescent theory. Handbook of Statistical Genomics: Two Volume Set pp. 145–30 (2019)

[35] Olafsson, S., Anderson, C.A.: Somatic mutations provide important and unique insights into the biology of complex diseases. Trends in Genetics (2021)

[36] Posada, D.: Cellcoal: coalescent simulation of single-cell sequencing samples. Molecular biology and evolution 37(5), 1535–1542 (2020)

[37] Rajaraman, A., Ullman, J.D.: Mining of massive datasets. Cambridge University Press (2011)

[38] Rausch, T., Zichner, T., Schlattl, A., Stütz, A.M., Benes, V., Korbel, J.O.: Delly: structural variant discovery by integrated paired-end and split-read analysis. Bioinformatics 28(18), i333–i339 (2012)

[39] Salk, J.J., Fox, E.J., Loeb, L.A.: Mutational heterogeneity in human cancers: origin and consequences. Annual Review of Pathology: Mechanisms of Disease 5, 51–75 (2010)

[40] Schmitt, M.W., Kennedy, S.R., Salk, J.J., Fox, E.J., Hiatt, J.B., Loeb, L.A.: Detection of ultra-rare mutations by next-generation sequencing. Proceedings of the National Academy of Sciences 109(36), 14508–14513 (2012). https://doi.org/10.1073/pnas.1208715109, https://www.pnas.org/content/109/36/14508

[41] Slaney, M., Casey, M.: Locality-sensitive hashing for finding nearest neighbors [lecture notes]. IEEE Signal processing magazine 25(2), 128–131 (2008)

[42] Slatko, B.E., Gardner, A.F., Ausubel, F.M.: Overview of nextgeneration sequencing technologies. Current protocols in molecular biology 122(1), e59 (2018)

[43] Stratton, M.R., Campbell, P.J., Futreal, P.A.: The cancer genome. Nature 458(7239), 719–724 (2009)

[44] Sudmant, P.H., Rausch, T., Gardner, E.J., Handsaker, R.E., Abyzov, A., Huddleston, J., Zhang, Y., Ye, K., Jun, G., Fritz, M.H.Y., et al.: An integrated map of structural variation in 2,504 human genomes. Nature 526(7571), 75–81 (2015)

[45] Voronina, N., Wong, J.K., Hübschmann, D., Hlevnjak, M., Uhrig, S., Heilig, C.E., Horak, P., Kreutzfeldt, S., Mock, A., Stenzinger, A., et al.: The landscape of chromothripsis across adult cancer types. Nature communications 11(1), 1–13 (2020)

[46] Wijewardhane, N., Dressler, L., Ciccarelli, F.D.: Normal somatic mutations in cancer transformation. Cancer Cell 39(2), 125–129 (2021)

[47] Yang, H., Lu, B., Lai, L.H., Lim, A.H., Alvarez, J.J.S., Zhai, W.: Psite: a phylogeny guided simulator for tumor evolution. Bioinformatics 35(17), 3148–3150 (2019)

[48] Zeira, R., Shamir, R.: Sorting cancer karyotypes using doublecut-and-joins, duplications and deletions. Bioinformatics 37(11), 1489–1496 (2021)

[49] Zhao, M., Liu, D., Qu, H.: Systematic review of next-generation sequencing simulators: computational tools, features and perspectives. Briefings in functional genomics 16(3), 121–128 (2017)

